# RGS14 is neuroprotective against seizure-induced mitochondrial oxidative stress and pathology in hippocampus

**DOI:** 10.1101/2023.02.01.526349

**Authors:** Nicholas H Harbin, Daniel J Lustberg, Cheyenne Hurst, Jean-Francois Pare, Kathryn M Crotty, Alaina L Waters, Samantha M Yeligar, Yoland Smith, Nicholas T Seyfried, David Weinshenker, John R Hepler

**Affiliations:** Emory University School of Medicine

**Keywords:** RGS14, hippocampus, CA2, kainic acid, seizure, oxidative stress, mitochondria, neuronal injury, microglia

## Abstract

RGS14 is a complex multifunctional scaffolding protein that is highly enriched within pyramidal cells (PCs) of hippocampal area CA2. There, RGS14 suppresses glutamate-induced calcium influx and related G protein and ERK signaling in dendritic spines to restrain postsynaptic signaling and plasticity. Previous findings show that, unlike PCs of hippocampal areas CA1 and CA3, CA2 PCs are resistant to a number of neurological insults, including degeneration caused by temporal lobe epilepsy (TLE). While RGS14 is protective against peripheral injury, similar roles for RGS14 during pathological injury in hippocampus remain unexplored. Recent studies show that area CA2 modulates hippocampal excitability, generates epileptiform activity and promotes hippocampal pathology in animal models and patients with TLE. Because RGS14 suppresses CA2 excitability and signaling, we hypothesized that RGS14 would moderate seizure behavior and early hippocampal pathology following seizure activity. Using kainic acid (KA) to induce status epilepticus (KA-SE) in mice, we show loss of RGS14 (RGS14 KO) accelerated onset of limbic motor seizures and mortality compared to wild type (WT) mice, and that KA-SE upregulated RGS14 protein expression in CA2 and CA1 PCs of WT. Utilizing proteomics, we saw loss of RGS14 impacted the expression of a number of proteins at baseline and after KA-SE, many of which associated unexpectedly with mitochondrial function and oxidative stress. RGS14 was shown to localize to the mitochondria in CA2 PCs of mice and reduce mitochondrial respiration *in vitro*. As a readout of oxidative stress, we found RGS14 KO dramatically increased 3-nitrotyrosine levels in CA2 PCs, which was greatly exacerbated following KA-SE and correlated with a lack of superoxide dismutase 2 (SOD2) induction. Assessing for hallmarks of seizure pathology in RGS14 KO, we observed worse neuronal injury in area CA3 (but none in CA2 or CA1), and a lack of microgliosis in CA1 and CA2 compared to WT. Together, our data demonstrates a newly appreciated neuroprotective role for RGS14 against intense seizure activity in hippocampus. Our findings are consistent with a model where, after seizure, RGS14 is upregulated to support mitochondrial function and prevent oxidative stress in CA2 PCs, limit seizure onset and hippocampal neuronal injury, and promote microglial activation in hippocampus.

## 1. Introduction

Regulators of G protein signaling (RGS) proteins are important negative modulators of G protein-dependent signaling, especially in the brain, where many are selectively and dynamically expressed in distinct brain regions to affect synaptic transmission, neuronal excitability, plasticity, and survival with consequences on behavior and disease (Gerber et al., 2016; Gold et al., 1997; Larminie et al., 2004). One such RGS protein, RGS14, is highly expressed in area CA2 pyramidal cells (PCs) of the hippocampus, the entorhinal and piriform cortex, basal ganglia and amygdala (Evans et al., 2014; Foster et al., 2021; Lee et al., 2010; Squires et al., 2018; Squires et al., 2021)). Behavioral consequences of global RGS14 deletion in mice (RGS14 KO) include enhancement of spatial learning (Lee et al., 2010), sensitization to cued fear learning (Alexander et al., 2019), and locomotor sensitivity to cocaine (Foster et al., 2021). Apart from these behavioral findings, insights into RGS14 function come largely from cellular studies assessing RGS14 interacting partners and effects on cell signaling. In addition to its canonical interactions with Gαi/o proteins that limit Gα activation (Hollinger et al., 2001), RGS14 is a multifunctional scaffold that possesses capacity to interact with many other signaling proteins, including activated monomeric GTPases (H-Ras, Rap2) and RAFkinases that modulate MAPK/ERK signaling (Shu et al., 2010; Traver et al., 2000; Vellano et al., 2013), calcium-dependent calmodulin (Ca^++^/CaM) and its effector kinase CaMKII (Evans et al., 2018a), 14-3-3γ (Gerber et al., 2018), and the NHERF1 PDZ scaffold (Friedman et al., 2022) among others (Harbin et al., 2021). RGS14 interaction with its binding partners is driven, in part, by GPCR activation (Vellano et al., 2013; Vellano et al., 2011). Unusual for a large multidomain protein, RGS14 shuttles between plasma membrane, cytosolic, and nuclear compartments (Branch and Hepler, 2017; Cho et al., 2005; Squires et al., 2021), implicating its importance and complexity in brain physiology, which has been relatively understudied compared to its roles in cell signaling.

The hippocampus is the major focus of temporal lobe-induced seizures and is susceptible to cell damage and injury (Blumcke et al., 2013). Of note, hippocampal area CA2, where RGS14 is expressed, is highly resistant to seizure-induced neuronal injury (Hatanpaa et al., 2014; Steve et al., 2014). In CA2 hippocampal physiology, RGS14 reduces intrinsic CA2 PC excitability and suppresses synaptic potentiation and structural plasticity at CA3-CA2 synapses (Evans et al., 2018b; Lee et al., 2010). This is in stark contrast to CA3-CA1 synapses that robustly express plasticity. RGS14 suppresses glutamate-induced calcium transients and modulates G protein, ERK, and calcium-dependent signaling in CA2 PCs to limit plasticity at CA3-CA2 synapses (Evans et al., 2018b; Lee et al., 2010). Increased excitatory input from CA3 and enhanced excitability in CA2 may therefore result in a hyperactive CA2 in RGS14 KO mice. In human and animal models of temporal lobe epilepsy (TLE), the hippocampus is subjected to neuropathological changes including hyperexcitability, neuroinflammation, and neurodegeneration. Importantly, CA2 has been shown to modulate excitatory/inhibitory balance of the hippocampus (Boehringer et al., 2017) and contribute to hippocampal epileptiform activity in human TLE and mouse models of TLE (Kilias et al., 2022; Whitebirch et al., 2022). Recent reports show CA2 becomes enlarged and receives sprouting mossy fiber connections (another hallmark of epilepsy) in mouse models of SE and patients with TLE (Freiman et al., 2021; Haussler et al., 2016), leading to the hypothesis that CA2 survives to maintain the hippocampal network while also promoting the development of TLE. Most recently, a compelling report demonstrated enhanced CA2 activity and its contribution to spontaneous seizure generation in a mouse model of TLE (Whitebirch et al., 2022). Based on these observations, we proposed that RGS14 in CA2 PCs may be protective and/or modulate CA2 activity during seizure by influencing excitation and/or intracellular signaling. Consistent with this idea, mice lacking RGS14 suffer worse injury and inflammation at the sites of injury in animal models of cardiac hypertrophy and hepatic ischemic-reperfusion injury (Li et al., 2016; Zhang et al., 2022).

Given that RGS14 is protective to insult injury in peripheral tissues, and is highly expressed in injury-resistant hippocampal area CA2, we set out to determine whether loss of RGS14 (RGS14 KO) would impact seizure behavior, seizure-induced cell injury and exacerbate hippocampal pathology following KA-SE. Additionally, we sought to determine how RGS14 may be affording this protection. Here, we report that RGS14 protein is upregualated in hippocampus following KA-SE induction, and that loss of RGS14 accelerated the onset of behavioral seizures and mortality during seizure. Using proteomic approaches, we determined changes in the hippocampal proteome in RGS14 KO animals, and identified altered metabolic/mitochondrial protein expression in RGS14 KO mice after seizure activity. Further investigation identified RGS14 localization to mitochondria in area CA2, and we provide evidence that RGS14 suppresses mitochondrial oxygen consumption *in vitro*. Consistent with this function, we show that mice lacking RGS14 exhibit enhanced oxidative stress, elevated neuronal injury, and altered inflammation following KA-induced seizure in vivo. Together, these finding suggest that RGS14 serves a previously unappreciated protective role in hippocampal seizure pathology

## 2. Materials and Methods

### 2.1. Animal Care

Male and female C57BL/6J wild-type (WT) and RGS14 null (RGS14 KO) mice ages 3-7 months were used in this study. Animals in all experiments were housed under a 12 h/12 h light/dark cycle with access to food and water ad libitum. All experimental procedures conformed to US NIH guidelines and were approved by the animal care and use committees (IACUC) of Emory University.

### 2.2. Kainic acid-induction of status epilepticus and monitoring of status epilepticus

Seizure-induction protocols using kainic acid (KA) were followed as described (Rojas et al., 2014). Wild type (WT) and RGS14 KO mice were individually housed in clean mouse cages without access to food and water 30 minutes prior to KA injection. KA (3 mg/mL) was prepared on the day of experimentation by dissolving KA (Tocris, 0222) in a 0.9% bacteriostatic saline solution. Mice were injected intraperitoneally (i.p.) with a single, 30 mg/kg dose of KA or 0.3 mL of saline (control). Immediately after injection, mice were monitored and scored for seizure behavior using a modified Racine scale (Racine, 1972; Rojas et al., 2014). Each mouse was given a behavioral seizure score from a scale of 0-6 every 5 minutes for 90 minutes. Mice were scored based off the most severe seizure behavior that was observed during the 5 minute interval. When an mouse reached mortality, they were given a score of 7, which was not included in the mean behavioral seizure score and no further scores were given for those mice. Ninety minutes after injection, all mice were injected intraperitoneally with 10 mg/kg diazepam to terminate seizure activity. After seizure termination, mice remained singly housed, and food (dry and moistened) and water were returned to the cage, and monitored for well-being.

### 2.3. Hippocampal tissue collection

To collect hippocampal sections for histology, male and female mice were anesthetized with sodium pentobarbital (Fatal Plus, 380 mg/kg, i.p.) and euthanized by transcardial perfusion with 4% paraformaldehyde (PFA) in PBS one day after KA or saline injection. After decapitation, mouse brains were dissected and post-fixed in 4% PFA at 4°C for 24 hours. After post-fix, brains were transferred to a 30% sucrose in PBS solution for 72 hours. Mouse brains were embedded in optimal cutting temperature (OCT) medium (Fisher Healthcare, 23730571), and coronal sections of hippocampus were collected at a thickness of 40 μm on a cryostat. Hippocampal sections were stored in 0.02% sodium azide in PBS at 4°C until they were evaluated for neuropathology. For proteomics and Western Immunoblot, mice were anesthetized with isoflurane followed by rapid decapitation and dissection of whole hippocampi. Whole hippocampi were snap frozen in liquid nitrogen and stored at −80°C until homogenization.

### 2.4. Western Immunoblotting

Frozen hippocampi were thawed on ice and homogenized in a modified RIPA buffer [150 mM NaCl, 50 mM Tris, 1 mM EDTA, 2 mM DTT, 5 mM MgCl_2_, 1X protease inhibitor (Roche, A32955) 1X Halt phosphatase inhibitor (ThermoFisher, 78428)]. Cells were lysed in modified RIPA buffer for 1 hour at 4°C, end-over-end. Lysates were cleared, and proteins were denatured by boiling in Laemmli buffer for 5 minutes (stored at −20°C until immunoblotting). Samples were resolved on 4-20% Mini-PROTEAN^®^ TGX™ precast protein gels (BioRad, 4561094), and gels were UV-activated for 45 seconds for total protein illumination (BioRad Stain-Free Technology) (Gilda and Gomes, 2015). Samples were then transferred to nitrocellulose membranes, and total protein was detected by activating membranes with ultraviolet light. Membranes were incubated in blocking buffer (5% non-fat milk in TBS containing 0.1% Tween-20 and 0.02% sodium azide) for 1 hour at room temperature. Membranes were incubated with anti-RGS14 (Neuromab, 75-170; 1:500) diluted in blocking buffer at 4°C overnight. Membranes were washed thrice with TBS containing 0.1% Tween-20 followed by incubation with goat anti-mouse IgG HRP-conjugate (Jackson ImmunoResearch, 115-035-003; 1:5000). Membranes were washed thrice with TBS containing 0.1% Tween-20, and enhanced chemiluminescence was used to develop the blots. Blots were imaged using a ChemiDoc MP Imager (BioRad). ImageLab (BioRad) software was used for densitometry analysis of RGS14 bands, and quantified band density was normalized to total protein content. Normalized RGS14 band densities are expressed relative to the mean of the saline group.

### 2.5. Immunohistochemistry

To determine protein expression levels in tissue by immunohistochemistry (IHC), free floating hippocampal sections from KA- and saline-treated WT and RGS14 KO mice were subjected to antigen retrieval by boiling sections in 10 mM citrate buffer for 3 minutes and blocked in 5% normal goat serum in 0.1% Triton-PBS (blocking solution) for 1 hour at room temperature. Sections were incubated overnight at 4°C with primary antibody specific for RGS14 (Neuromab, 75-170, 1:500, WT sections only), IBA1 (Wako, 019-19741, 1:1000), 3-nitrotyrosine (Abcam, ab61392, 1:500), or SOD2 (Proteintech, 24127-1-AP, 1:1000) diluted in blocking solution. Sections were washed thrice with PBS prior to incubation with the appropriate fluorescent secondary. Goat anti-mouse Alexa Fluor Plus 488 (ThermoFisher, A32723, 1:500), goat anti-rabbit Alexa Fluor 546 (ThermoFisher, A-11010, 1:500), or NeuroTrace™ 660 Nissl (ThermoFisher, N21483, 1:500) were diluted in blocking solution, and sections were blocked from light and incubated in secondary antibody solution for 2 hours at room temperature. Sections were washed thrice with PBS and mounted onto SuperFrost Plus microscope slides (Thermo Fisher Scientific). Once dried, sections were coverslipped with DAPI Fluoromount G (Southern Biotech, 0100-20) and stored away from light at 4°C prior to imaging.

### 2.6. Fluorescent Imaging and quantification of immunofluorescence

Fluorescent micrographs were collected from dorsal hippocampal sections using a Leica DM6000B epifluorescent upright microscope at 5X, 10X, and 20X magnifications using the appropriate wavelength filters. Images magnified at 20X were processed, analyzed, and quantified using ImageJ software. For RGS14 and SOD2, standardized regions of interests were drawn around each layer [stratum radiatum (SR), stratum pyramidale (SP), stratum oriens (SO)] of each CA field (CA1, CA2, CA3) and the dentate gyrus (DG). For 3-nitrotyrosine, a standardized ROI was drawn around each CA field incorporating the three layers (SR, SP, SO). Quantification for 3-nitrotyrosine was performed this way because 3-nitrotyrosine accumulation was consistent throughout all layers, while immunofluorescence of RGS14 and SOD2 expression was noticeably different depending on the layer of analysis. For quantification of raw immunofluorescence, background correction and intensity thresholding (Otsu method) were performed consistently across all images for each ROI prior to quantifying raw immunofluorescence.

### 2.7. Quantification of microglia cell counts and size

For the microglia marker ionized calcium binding adaptor molecule 1 (IBA1), a standardized region of interest was drawn for each CA subfield incorporating the stratum radiatum (SR), stratum pyramidale (SP), and stratum oriens (SO) and the DG incorporating the hilus, granule cell layer, and molecular layer. Background subtraction, intensity thresholding (Otsu method), and size and shape criteria for microglia (25-1000 μm^2^, circularity 0.05-1) were applied consistently across all images for each ROI using ImageJ software. Cell counts and area of IBA1 positive cells were quantified for each image. Microglia density is expressed as the number of cells divided by the area of ROI, and average size is expressed at the mean area of IBA1 positive cells.

### 2.8. Tissue homogenization and digestion of proteomics

Each tissue sample was added to 250 μL of urea lysis buffer (8 M urea, 10 mM Tris, 100 mM NaH2PO4, pH 8.5, including 2.5 μL (100x stock) HALT(-EDTA) protease and phosphatase inhibitor cocktail (Pierce)) in a 1.5 mL Rino tube (Next Advance) harboring stainless steel beads (0.9-2 mm in diameter). Samples were homogenized twice for 5-minute intervals in the cold room (4 °C). Protein homogenates were transferred to 1.5 mL Eppendorf tubes on ice and were sonicated (Sonic Dismembrator, Fisher Scientific) thrice for 5 sec each with 5 sec intervals of rest at 30% amplitude to disrupt nucleic acids and were subsequently centrifuged at 4° C. Protein concentration was determined by the bicinchoninic acid (BCA) method, and samples were frozen in aliquots at −80 °C. Protein homogenates (100 μg) were treated with 1 mM dithiothreitol (DTT) at room temperature for 30 min, followed by 5 mM iodoacetimide at room temperature for 30 min in the dark. Protein samples were digested with 1:25 (w/w) lysyl endopeptidase (Wako) at room temperature overnight. Next day, samples were diluted with 50 mM NH4HCO3 to a final concentration of less than 2 M urea and were further digested overnight with 1:25 (w/w) trypsin (Promega) at room temperature. The resulting peptides were desalted with HLB column (Waters) and were dried under vacuum.

### 2.9. Liquid chromatography coupled to tandem spectrometry (LC-MS/MS)

The data acquisition by LC-MS/MS was adapted from a published procedure (Seyfried et al., 2017). Derived peptides were resuspended in the loading buffer (0.1% trifluoroacetic acid, TFA) and were separated on a Water’s Charged Surface Hybrid (CSH) column (150 μm internal diameter (ID) x 15 cm; particle size: 1.7 μm). The samples were run on an EVOSEP liquid chromatography system using the 15 samples per day preset gradient (88 min) and were monitored on a Q-Exactive Plus Hybrid Quadrupole-Orbitrap Mass Spectrometer (ThermoFisher Scientific). The mass spectrometer cycle was programmed to collect one full MS scan followed by 20 data dependent MS/MS scans. The MS scans (400-1600 m/z range, 3 x 106 AGC target, 100 ms maximum ion time) were collected at a resolution of 70,000 at m/z 200 in profile mode. The HCD MS/MS spectra (1.6 m/z isolation width, 28% collision energy, 1 x 105 AGC target, 100 ms maximum ion time) were acquired at a resolution of 17,500 at m/z 200. Dynamic exclusion was set to exclude previously sequenced precursor ions for 30 seconds. Precursor ions with +1, and +7, +8 or higher charge states were excluded from sequencing.

### 2.10. Label-free quantification (LFQ) using MaxQuant

Label-free quantification analysis was adapted from a published procedure (Seyfried et al., 2017). Spectra were searched using the search engine Andromeda, integrated into MaxQuant, against mouse database (86,470 target sequences). Methionine oxidation (+15.9949 Da), asparagine and glutamine deamidation (+0.9840 Da), and protein N-terminal acetylation (+42.0106 Da) were variable modifications (up to 5 allowed per peptide); cysteine was assigned as a fixed carbamidomethyl modification (+57.0215 Da). Only fully tryptic peptides were considered with up to 2 missed cleavages in the database search. A precursor mass tolerance of ±20 ppm was applied prior to mass accuracy calibration and ±4.5 ppm after internal MaxQuant calibration. Other search settings included a maximum peptide mass of 6,000 Da, a minimum peptide length of 6 residues, 0.05 Da tolerance for orbitrap and 0.6 Da tolerance for ion trap MS/MS scans. The false discovery rate (FDR) for peptide spectral matches, proteins, and site decoy fraction were all set to 1 percent. Quantification settings were as follows: re-quantify with a second peak finding attempt after protein identification has completed; match MS1 peaks between runs; a 0.7 min retention time match window was used after an alignment function was found with a 20-minute RT search space. Quantitation of proteins was performed using summed peptide intensities given by MaxQuant. The quantitation method only considered razor plus unique peptides for protein level quantitation.

### 2.11. Differential Expression and Gene Ontology (GO) Analysis

LFQ normalized abundances summarized for protein level quantification based on parsimoniously assembled razor plus unique peptides in MaxQuant were considered for statistical comparisons between WT and RGS14 KO sample abundances after either SAL or KA. Prior to statistical comparison, network connectivity was used to identify and remove sample outliers, which were defined as samples of 3 or more standard deviations away from the mean. In addition, only proteins with quantitation in more than 50% of the samples were included and log2 transformed. Significantly differentially changed proteins between groups were defined used one-way ANOVA with Tukey’s comparison post hoc test. Differential expression displayed as volcano plots were generated using the GraphPad Prism. Gene ontology (GO), Wikipathway, Reactome and molecular signatures database (MSigDB) term enrichment in our gene sets of mouse network module members and significantly differentially changed protein subsets was determined by gene set enrichment analysis (GSEA) using an in-house developed R script (https://github.com/edammer/GOparallel). Briefly, this script performs one tailed Fisher’s exact tests (FET) enabled by functions of the R piano package for ontology enrichment analysis on gene sets downloaded from http://baderlab.org/GeneSets, which is maintained and updated monthly to pull in current gene sets from more than 10 different database sources including those mentioned above (Reimand et al., 2019; Varemo et al., 2013). Redundant core GO terms were pruned in the GOparallel function using the minimal_set function of the ontologyIndex R package (McKenzie et al., 2017). All proteomic statistical analyses were performed in R (version 4.0.3).

### 2.12. Immunogold electron microscopy

Mouse brain tissue sections from wild type animals containing the dorsal hippocampus were processed for a pre-embedding immunogold procedure to characterize subcellular RGS14 expression in area CA2 as previously performed (Squires et al., 2018). The sections were pre-incubated in a 1% sodium borohydride/PBS solution for 20 min and washed in PBS. Sections were then put in a cryoprotectant solution for 20 minutes before being frozen at −20°C. Sections were then thawed and returned to a graded series of cryoprotectant and PBS. This was followed by an incubation in 5% milk diluted in PBS for 30 min and then and overnight incubation at room temperature in primary RGS14 antibody solution consisting of RGS14 antibody (rabbit pAb – Protein Tech, 16258-1-AP; 1:4000 dilution) and 1% dry milk in TBS-gelatin buffer (0.02 M, 0.1% gelatin, pH 7.6). After this incubation, sections were rinsed in TBS-gelatin and incubated for 2 hours with secondary goat anti-rabbit Fab fragments conjugated to 1.4-nm gold particles (1:100; Nanoprobes, Yaphank, NY) and 1% dry milk in TBS-gelatin to limit cross-reactivity of the secondary antibody. To optimize RGS14 visualization, sections underwent incubation for approximately 10 min in the dark with a HQ Silver Kit (Nanoprobes) to increase gold particle sizes to 30–50 nm through silver intensification. The sections were then rinsed in phosphate buffer (0.1 M, pH 7.4) and treated with 0.5% osmium for 10 min and 1% uranyl acetate in 70% ethanol for 10 min, followed by dehydration with increasing ethanol concentrations. They were then placed in propylene oxide, embedded in epoxy resin (Durcupan ACM, Fluka, Buchs, Switzerland) for at least 12 hours, and baked in a 60 °C oven for 48 hours. As controls, sections were incubated in a solution devoid of the RGS14 antibody but containing the secondary antibody and the rest of the immunostaining protocol remained the same as above.

### 2.13. Measurement of Mitochondrial Respiration using Seahorse XF Cell Mitochondrial Stress Test in HEK293T cells

The measurement of mitochondrial respiration was performed as previously described with modifications (Morris et al., 2021). Briefly, HEK293T cells were cultured in Dulbecco’s essential medium (DMEM; Gibco, 11-995-040) supplemented with 10% fetal bovine serum and 1% penicillin/streptomycin and incubated at 37°C, 5% CO_2_ until the time of seeding or passaging. Cells were maintained by passaging every 2-3 days when cells reached 70-80% confluency. Mitochondrial respiration was evaluated using a XFe96 Extracellular Flux Analyzer (Agilent Seahorse Bioscience Inc., Billerica, MA) and Seahorse CF Cell Mito Stress Test Kit (Agilent, 103015-100) according to manufacturer’s instructions. Briefly, cells were seeded into a Seahorse XFe96 microplate (Agilent, 103794-100) at 15,000 cells/well and incubated for 24 hr at 37°C, 5% CO_2_. FLAG-RGS14 or pcDNA3.1 (negative control) plasmids (Shu et al., 2007) were transiently transfected using transfection medium DMEM supplemented with 5% FBS and 1% penicillin/streptomycin and polyethyleneimine (PEI) as the transfection reagent. Cells were then incubated for 24 hr at 37°C, 5% CO_2_ to ensure adequate expression of both constructs. After transfection, cells were switched to Seahorse XF Base Medium supplemented with 1mM L-glutamine, 5.5 mM D-glucose, and 2 mM sodium pyruvate (pH of 7.4) and equilibrated in this medium for 30 minutes. Oxygen consumption rate (OCR) was measured prior to and after sequential treatment with 1 μM oligomycin (mitochondrial complex V inhibitor), 0.5 μM carbonilcyanide p-trifluoromethoxyphenylhydrazone (FCCP) (ATP synthase inhibitor and proton uncoupler), and 0.5 μM rotenone/antimycin A (Complex I/III inhibitor). Basal respiration, mitochondrial ATP-linked respiration, maximal respiration, proton leak, spare respiratory capacity, and non-mitochondrial linked respiration were determined using the XF Wave 2.1 software. Cell lysates were collected in lysis buffer, and Pierce BCA assay was used to determine protein concentration. OCR values were normalized to HEK293T protein concentration in the same sample and were expressed as mean ± SEM. Western immunoblotting to confirm protein expression was performed as stated above using anti-FLAG HRP-conjugated antibody (Sigma Aldrich, A8592; 1:15,000) to verify transfection and expression of FLAG-RGS14.

### 2.14. FluoroJade B (FJB) tissue staining

To assess for neurodegeneration one day after KA-SE, FluoroJade B (FJB) (Histochem, #1FJB, Jefferson, AR) was used to label and quantify degenerating neurons (Schmued et al., 1997) according to manufacturer’s instructions with modifications (Rojas et al., 2014). Briefly, hippocampal sections were mounted and dried on SuperFrost Plus microscope slides. Once dried, sections were sequentially immersed with 80% EtOH, 70% EtOH, and ddH2O for 2 minutes each. Sections were incubated with 0.06% KMnO4 (dissolved in ddH2O) for 10 minutes and rinsed twice with ddH2O for 2 minutes each to remove all traces of KMnO4. Sections were incubated in a 0.0004% solution of Fluoro-Jade B for 30 minutes covered from light. Sections were then rinsed thrice with ddH2O for 1 minute each and air dried for 24 hours while covered from light. Sections were cleared with xylenes for 5 minutes and coverslipped with DPX (Electron Microscopy Sciences, Hatfield, PA) mounting medium. FJB labeled neurons were visualized and imaged using a Leica DM6000B epifluorescent upright microscope with filters for fluorescein or FITC at 10X and 20X magnifications. Representative images were collected from dorsal hippocampal CA fields. A standardized region of interest (ROI) was drawn around CA3 cell bodies in the stratum pyramidale (SP) from 20X magnified images. Fluoro-Jade B positive cells were quantified using DotDotGoose (Ersts, American Museum of Natural History, Center of Biodiversity and Conservation, version 1.5.3). Cell counts are represented as the total number of Fluoro-Jade B positive cells in each representative section for each animal.

### 2.15. Statistical analysis

All statistical analysis was performed in GraphPad Prism (version 9.3.1). Data were subjected to Grubb’s outlier analysis prior to statistical comparisons. For behavioral seizure experiments, Gehan-Breslow-Wilcoxon test was used to determine differences between survival curves, and Fisher’s exact test was used to determine contingency of genotype on binned mortality and overall mortality. Multiple t-tests were used to compare mean behavioral seizure score at each time point during the 90 minute behavioral seizure period, and unpaired Student’s t-test was used to compare mean latency to Stage 3 seizure activity between genotypes. Two-way ANOVA with Sidak post-hoc comparison was used to assess effects of genotype and sex on latency to mortality and Stage 3 seizure activity. One-way ANOVA with Dunnett’s post-hoc comparisons was used to compare mean RGS14 expression (band density from immunoblot) in hippocampal lysates, and unpaired Student’s t-tests were used to compare mean RGS14 immunofluorescence between SAL and 1d KA treatments. Unpaired Students t-tests were used to compare OCR means between control and RGS14 overexpression groups, SOD2 mean immunofluorescence in CA1, and mean number of FJB+ neurons in CA3 between WT and RGS14 KO mice after KA-SE. For comparison 3-NT and SOD2 mean immunofluorescence and IBA1+ microglia cell density and mean area, two-way ANOVA with Tukey’s or Sidak’s post-hoc analysis was used.

## 3. Results

### 3.1 Loss of RGS14 increases susceptibility to KA-induced seizures by expediting entry into SE and mortality

Recent evidence has supported a role for hippocampal area CA2 in modulating hippocampal network activity and seizure generation (Boehringer et al., 2017; Kilias et al., 2022; Whitebirch et al., 2022). Because RGS14 is highly and selectively expressed within area CA2 where it reduces neuronal excitability and inhibits long-term potentiation at CA3-CA2 excitatory synapses (Lee et al., 2010), we explored whether loss of RGS14 (RGS14 KO) altered hippocampal seizure activity. To test this, we followed previously described seizure-induction protocols (Rojas et al., 2014). Specifically, we administered a single dose of 30 mg/kg KA by intraperitoneal (i.p.) injection into WT (n = 39 total; 26 males, 13 females) and RGS14 KO mice (n = 41 total; 25 males, 16 females) to induce status epilepticus (KA-SE). We monitored survival during seizure activity, progression of behavioral seizure severity, and latency to seizure staging for 90 minutes following KA administration. Behavioral seizure score (0-6) was measured every 5 minutes for 90 minutes using a modified Racine scale (see Materials and Methods and Table 1) until termination of seizure with diazepam (10 mg/kg, i.p.). Status epilepticus was defined as 30 minutes or more of continuous Stage 3 or higher seizure activity.

**Table 1.** Modified Racine scale to evaluate behavioral seizure progression in mice (1 column)

| Behavioral Score | Motor Behavior |
| --- | --- |
| **0 Normal Behavior** | walking, exploring, sniffing, grooming |
| **1 Freeze Behavior** | immobile, staring, rigged posture, hunched |
| **2 Automatisms** | head bobbing, face washing, whisker twitching, chewing, stargazing |
| **3 Early Seizure Behavior** | Myoclonic jerk, partial or whole body clonus |
| **4 Moderate Seizure Behavior** | rearing with bilateral forelimb clonus |
| **5 Advanced Seizure Behavior** | rearing and falling, loss of posture |
| **6 Intense Seizure Behavior** | tonic clonic seizure with running/bouncing |
| **7 Death** |  |

After KA administration, we observed that all mice regardless of genotype either entered SE (30 min or longer of > Stage 3 behavioral seizure activity) or died at higher stage seizures (data not shown). Comparison of survival curves between WT and RGS14 KO mice demonstrate that RGS14 KO mice die significantly quicker than WT mice (Fig. 1A), and the mean latency to mortality was significantly reduced in the RGS14 KO mice (Fig. 1B). At the 30-minute time interval, nearly half of RGS14 KO mice (49%, 19 of 41) reach mortality, while less than one-quarter of WT mice (23%, 6 of 39) reach mortality at the same interval (Fig. 1A). Of the mice that reach mortality, significantly more RGS14 KO mice (76%, 19 of 24) die in the first 30 minutes after KA administration compared to WT mice (33%, 6 of 18) (Fig. 1C). Surprisingly, there was no significant difference between the overall mortality rate between WT mice (46%, 18 of 39) and RGS14 KO mice (59%, 24 of 41) after the 90 minute seizure period (Fig. 1D).

**Figure 1.**
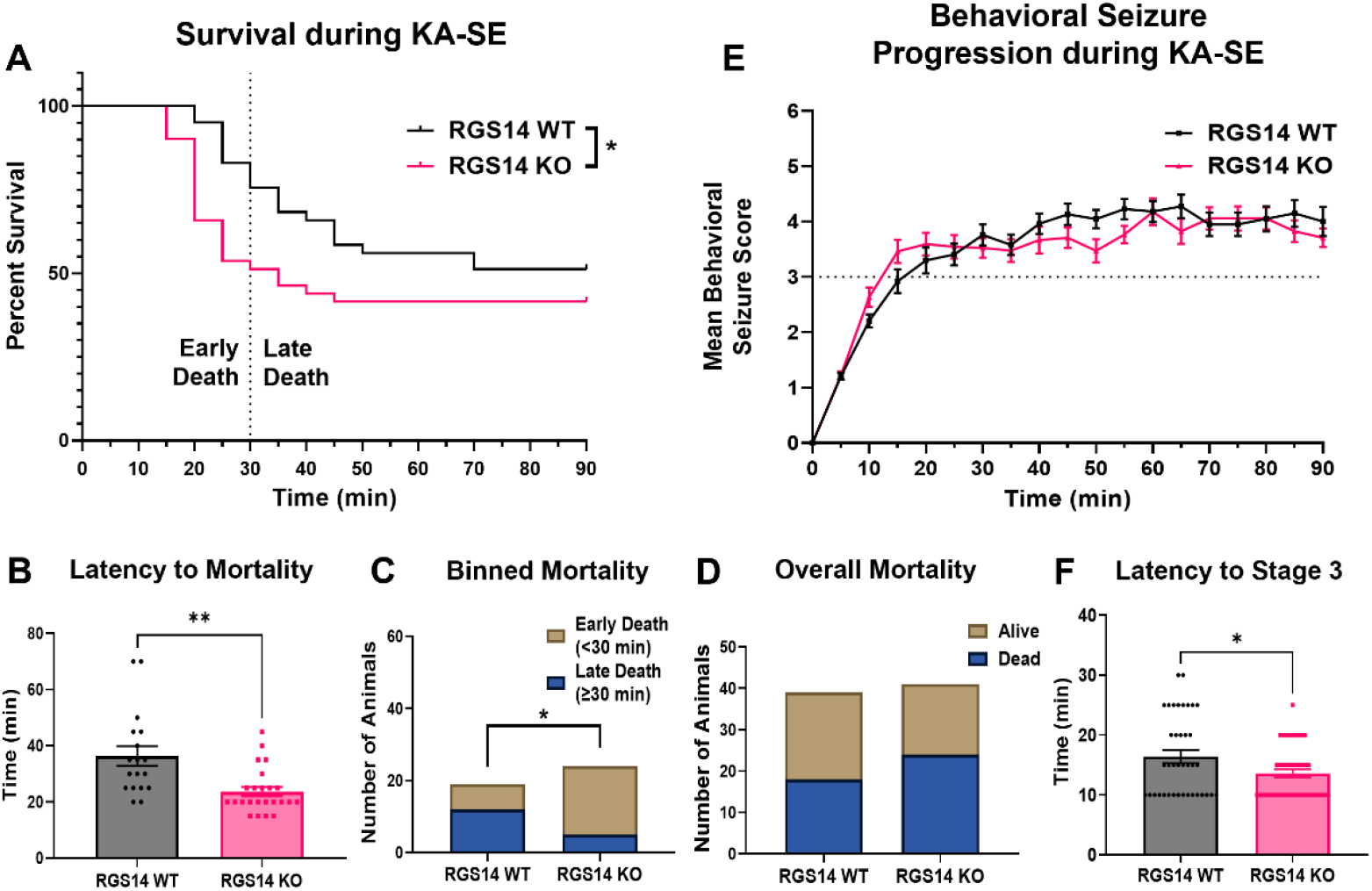
Loss of RGS14 increases susceptibility to KA-induced seizure by expediting mortality and entry into SE. (A) Survival curve comparing the percentage of alive animals between WT (n = 39) and RGS14 KO (n = 41) mice after 30 mg/kg KA injection shows RGS14 KO mice die quicker than WT mice following KA (KO/WT Hazard Ratio = 1.770, Mantel-Haenszel) (B) Mean latency to mortality in WT and RGS14 KO mice following KA injection (WT, 36.39 ± 3.52 min; RGS14 KO, 23.75 ± 1.63 min). (C) The number of animals that died in the first 30 minutes (Early Death; WT, n = 6; RGS14 KO, n = 19) or last 60 minutes (Late Death; WT, n = 12; RGS14 KO, n = 5) of the 90-minute seizure period. (D) The number of animals that were alive (WT, n = 21; RGS14 KO, n = 17) or dead (WT, n = 18; RGS14 KO, n = 24 dead) at the end of the 90-minute seizure period. (E) The mean behavioral seizure score plotted over time in WT (n = 39) and RGS14 KO (n = 41) mice. Seizure stage was scored every 5 minutes until seizure termination 90 minutes after KA injection. (F) The mean latency of animals to reach Stage 3 behavioral seizure activity after KA injection (WT, 16.41 ± 1.07 min; RGS14 KO, 13.54 ± 0.68 min). *Statistical analysis:* (A) Gehan-Breslow-Wilcoxon test to compare survival curves (*p < 0.05). (B, F) Unpaired t-test to compare latency to mortality (B; WT vs RGS14 KO, **p < 0.01) and Stage 3 seizure (F; WT vs RGS14 KO, *p < 0.05) (C, D) Fisher’s exact test to assess for contingency of genotype on binned morality (C, *p < 0.05) or overall mortality (D, p > 0.05). (E) Multiple t-tests were used to compare between genotypes at each time point (p > 0.05 for all time points). Error bars represent standard error of the mean (SEM). WT, wild-type; KA, kainic acid; SE, status epilepticus. (1.5 column)

Initial analysis of behavioral seizure scores demonstrated that WT and RGS14 KO mice progress similarly through behavioral seizure staging following KA (Fig. 1E). When latency to Stage 3 (i.e. SE) was compared between genotypes, we found RGS14 KO mice reached Stage 3 slightly but significantly quicker than WT mice (Fig. 1F). Because seizure susceptibility can differ between males and females, we assessed the contribution of sex to the seizure phenotype (Supplemental Fig. S1). We found only a main effect of genotype on the latency to mortality (Fig. S1A) and SE (Fig. S1B), where RGS14 KO had reduced latency to mortality in both males and females. These data suggest that RGS14 limits the behavioral seizure response to KA, where loss of RGS14 expedites entry into SE and mortality following KA without affecting overall mortality.

### 3.2. RGS14 protein expression is upregulated following status epilepticus

Several studies have demonstrated that RGS14 expression is significantly altered in animal and cellular models of cardiac hypertrophy and hepatic ischemic-reperfusion injury (Li et al., 2016; Zhang et al., 2022), and that the expression of other RGS proteins are regulated by seizure activity (Gold et al., 1997). Therefore, we sought to determine whether RGS14 expression is altered following KA-SE (Fig. 2). One, three, or seven days after KA-SE, whole hippocampi were collected from WT mice for protein analysis, and Western blot was used to detect RGS14 expression following in whole hippocampal lysate (Fig. 2A). Relative to saline-treated hippocampal tissue, there was a striking and significant upregulation of RGS14 protein following KA-SE. (Fig. 2B). Peaking one day after KA-SE, RGS14 expression is significantly induced more than 5-fold compared to saline (Fig. 2B). RGS14 expression remains high relative to saline three days after KA-SE but returns to near baseline levels 7 days after KA-SE (Fig. 2B). These findings suggest that RGS14 induction in the hippocampus is regulated by seizure activity and may play an important role in injury response following KA-SE.

**Figure 2.**
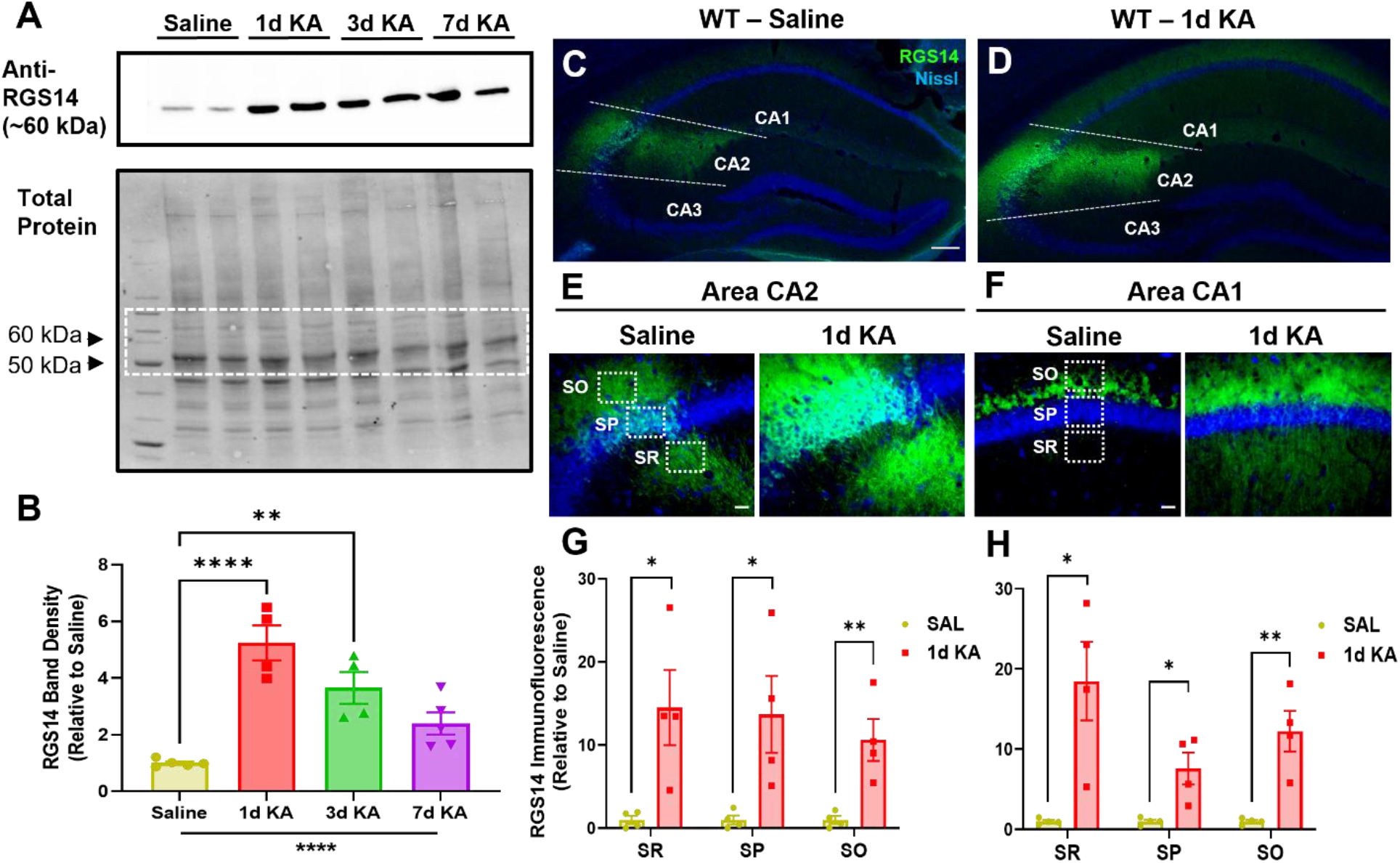
Hippocampal RGS14 protein expression is upregulated following KA-SE. (A) Representative Western blot showing RGS14 protein expression (top) relative to total protein (bottom) in WT hippocampal lysate 1 day after saline (n = 5) or 1 (n = 4), 3 (n = 4) or 7 (n = 5) days after KA-SE. Dashed box in total protein blot represents strip used for RGS14 immunoblot. (B) Quantification of mean RGS14 protein band density normalized to total protein density (saline, 1.00 ± 0.06; 1d KA, 5.24 ± 0.62; 3d KA, 3.65 ± 0.56; 7d KA, 2.40 ± 0.39) (C, D) Representative IHC images of RGS14 expression in the dorsal hippocampus of WT mice 1 day after saline (C) or KA-SE (D). Dashed lines divide the CA1, CA2, and CA3 subregions of the hippocampus. Scale bar, 100 μm. (E) Representative IHC image of area CA2 1 day after saline or KA-SE. (F) Representative IHC image of area CA1 1 day after saline or KA-SE. Dashed boxes illustrate regions of quantification for pyramidal cell layers (SR, SP, SO) in area CA2 (E) and CA1 (F). Scale bar, 20 μm. (G) Mean RGS14 immunofluorescence (n=4 per treatment) in saline- and KA-treated CA2 SR (saline, 1.00 ± 0.49; 1d KA, 14.52 ± 4.52), SP (saline, 1.00 ± 0.54; 1d KA, 13.69 ± 4.63), and SO (saline, 1.00 ± 0.47; 1d KA, 10.62 ± 2.53). (H) Mean RGS14 immunofluorescence (n=4 per treatment) in saline- and KA-treated CA1 SR (saline, 1.00 ± 0.19; 1d KA, 18.51 ± 4.91), SP (saline, 1.00 ± 0.22; 1d KA, 7.61 ± 1.99), and SO (saline, 1.00 ± 0.19; 1d KA, 12.26 ± 2.54). *Statistical Analysis:* (B) One-way ANOVA with Dunnett post-hoc comparisons to compare mean RGS14 band density (one-way ANOVA, F = 17.65 for treatment, ****p < 0.0001; Dunnet’s, 1d KA vs SAL, p < 0.0001; 3d KA vs SAL, p < 0.01; 7d KA vs SAL, p > 0.05). (G, H) Unpaired t-tests to compare mean RGS14 immunofluorescence in CA2 (SAL vs 1d KA: SR, SP, *p < 0.05; SO, **p < 0.01) and CA1 (SAL vs 1d KA: SR, SP, *p < 0.05; SO, **p < 0.01). p > 0.05 is considered not significant (ns). Error bars represent the SEM. IHC, immunohistochemistry; SR, stratum radiatum; SP, stratum pyramidale; SO, stratum oriens. (2 column)

To determine where in the hippocampus RGS14 expression is being induced, immunohistochemistry (IHC) was performed on dorsal hippocampal sections collected from WT mice one day after KA-SE or saline (SAL) (Fig. 2C-H). In saline-treated WT mice (Fig. 2C), RGS14 expression is mostly restricted to the cell bodies (*stratum pyramidale*, SP) and dendrites (*stratum radiatum*, SR, and *stratum oriens*, SO) of area CA2 PCs (Fig. 2E). Additionally, we observed modest RGS14 expression in area CA1, mostly in the SO of CA1 where CA2 axons project to CA1 pyramidal cell (Fig. 2F). One day following KA-SE (Fig. 2D), we detected a robust upregulation of RGS14 in CA2 PCs (Fig. 2E), where RGS14 expression is significantly induced throughout each layer of the region. Similarly, we observed upregulation of RGS14 expression in area CA1 one day after KA-SE (Fig. 2F). Like CA2, RGS14 upregulation was statistically significant in each layer of CA1 (Fig. 2H), particularly in the SR, where proximal dendrites of CA1 receive projections mostly from CA3, and the SO.

This demonstrates that RGS14 expression is robustly induced by KA-SE within hippocampal areas CA2 and CA1 (Fig. 2E, F), which likely reflects RGS14 upregulation in CA2 PC dendrites (CA2 SR, SO), cell bodies (CA2 SP), and axons (CA1 SO) (Fig. 2G) as well as CA1 PC dendrites (CA1 SR, SO) and cell bodies (CA1 SP) (Fig. 2H). Importantly, this is the first evidence demonstrating RGS14 expression is activity-dependent in the hippocampus, and suggests RGS14 may serve a protective role in this brain region following seizure.

### 3.3. Metabolic and mitochondrial proteins are differentially expressed between WT and RGS14 KO mice 1 day after KA-SE

To better understand how RGS14 may be influencing seizure activity in the hippocampus and the effects of losing RGS14 induction following seizure activity, we performed a proteomics analysis on hippocampal tissue taken from WT and RGS14 KO mice 1 day after saline or KA-SE treatment (n=5 per genotype and treatment) (Fig. 3 and Fig. 4). Whole hippocampi were homogenized, digested, and subjected to LC-MS/MS, and label-free quantitation (LFQ) was used to obtain relative protein abundance in each sample. Protein abundance was compared within treatment (SAL or KA), between genotype (WT vs RGS14 KO) to identify differentially expressed proteins (DEPs) in RGS14 KO hippocampi (Fig. 3A, B). For these analysis, 45 proteins in the SAL treatment and 49 proteins in the KA treatment were determined to be significantly up or downregulated between WT and RGS14 KO groups (adjusted p < 0.05), where 9 proteins were differentially expressed within both saline and KA treatments (Fig. 3C). These proteins are visualized in a volcano plot as either downregulated (left side) or upregulated (right side) in RGS14 KO hippocampi relative to WT hippocampi (Fig. 3A) (comprehensive table of DEPs listed in Supplementary Table 1).

**Figure 3.**
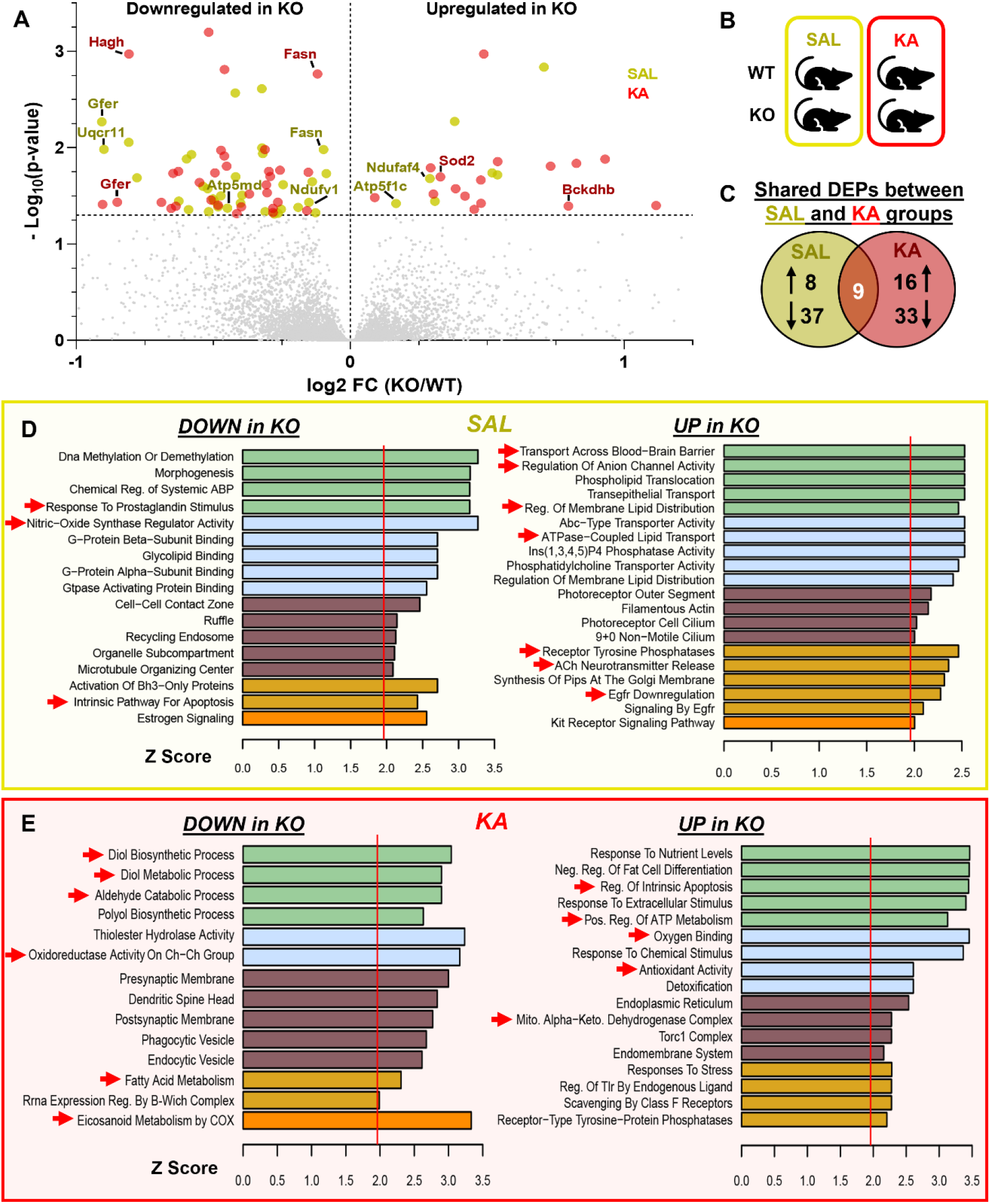
Differential expression analysis of the hippocampal proteome reveals altered mitochondrial and metabolic protein expression in RGS14 KO versus WT mice. (A) Volcano plot showing differentially expressed proteins (DEPs) one day after SAL (yellow) or KA-SE (red) treatment in RGS14 KO hippocampus relative to WT plotted by their −log_10_(p-value) and log_2_ fold change (KO/WT). Proteins were considered significantly downregulated if the adjusted p-value < 0.05, log_2_ fold change (KO/WT) < 0 and significantly upregulated if the adjusted p-value < 0.05, log_2_ fold change (KO/WT) > 0 (n = 5 per genotype/condition). Labeled dots represent DEPs involved with cellular metabolism and localize to the mitochondria. (B) Diagram illustrating between-genotype, within-treatment comparisons used to determine DEPs. (C) Venn diagram showing the number of upregulated or downregulated proteins in RGS14 KO relative to WT after SAL or KA-SE and DEPs common to both treatment groups. (D-E) Gene ontology analysis demonstrates similar biological function and processes of DEPs that are either downregulated (DOWN) or upregulated (UP) in RGS14 KO hippocampi one day after SAL (D) or KA-SE (E). *Red arrows* indicate ontologies of interest that may influence seizure susceptibility (D) or mitochondrial metabolism and oxidative stress (E). (2 column)

**Figure 4.**
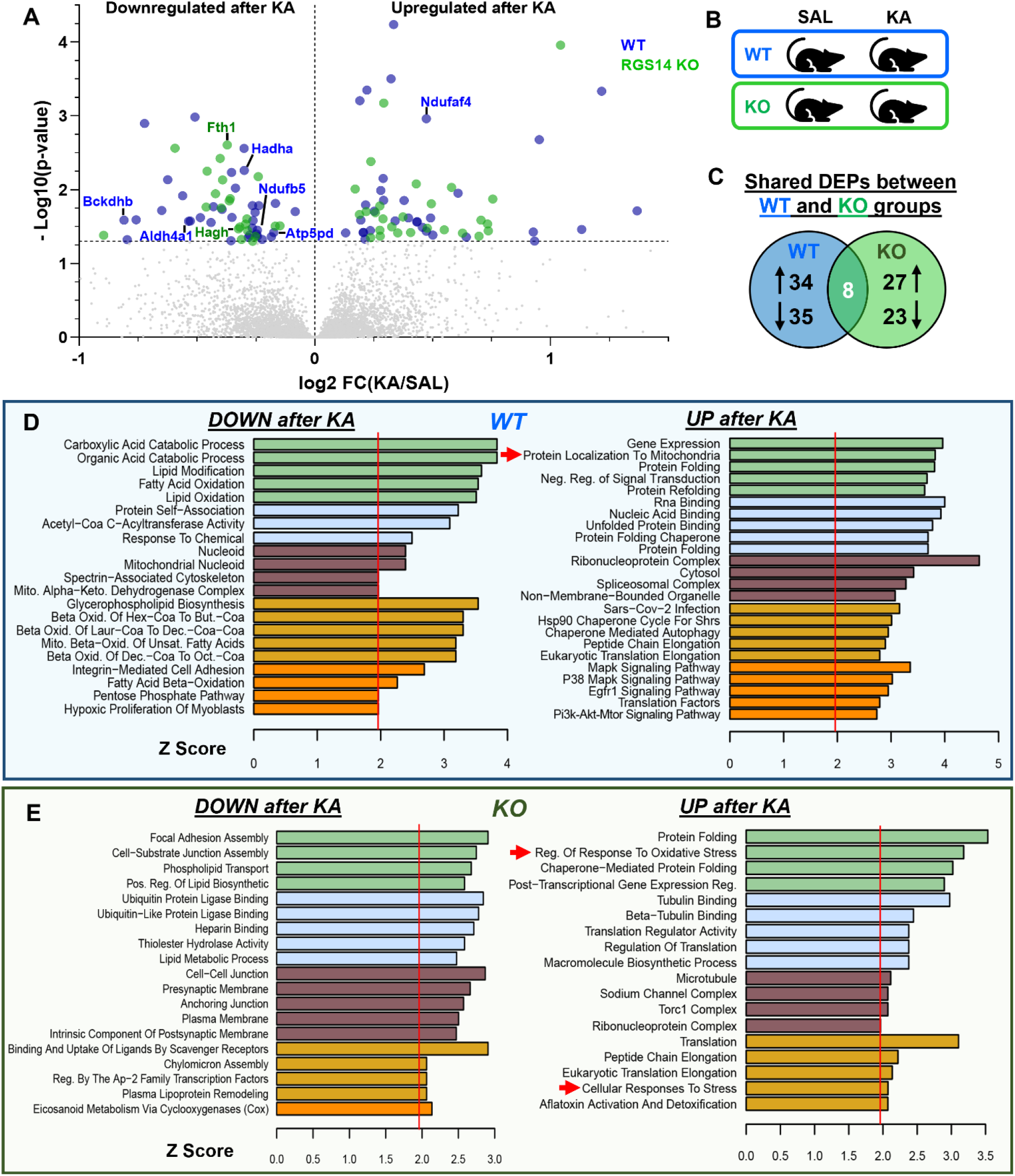
Comparison of WT and RGS14 KO hippocampal proteome after KA-SE suggests oxidative stress as a potential consequence of RGS14 KO. (A) Volcano plot showing differentially expressed proteins (DEPs) in WT or RGS14 KO one day after KA-SE plotted by their −log10(p-value) and log2 fold change (KA/SAL). Proteins were considered significantly downregulated if the adjusted p-value < 0.05, log_2_ fold change (KA/SAL) < 0 and significantly upregulated if the adjusted p-value < 0.05, log_2_ fold change (KA/SAL) > 0 (n = 5 per genotype/condition). Labeled dots represent DEPs involved with cellular metabolism and localize to the mitochondria. (B) Diagram illustrating within-genotype, between-treatment comparisons used to determine DEPs. (C) Venn diagram showing the number of upregulated or downregulated proteins following KA-SE in WT or RGS14 KO hippocampi and DEPs common to both genotypes. (D-E) Gene ontology analysis demonstrates similar biological function and processes of DEPs that are either downregulated (DOWN) or upregulated (UP) following KA-SE in WT (D) or RGS14 KO A (E). Red arrows indicate ontologies of interest related to metabolic and mitochondrial function. (2 column)

Gene ontology (GO) analysis was performed on up- and downregulated DEPs in SAL-treated mice identify common cellular pathways and processes associated with the DEPs that may influence sensitivity to KA (Fig. 3D). GO terms related to G-protein processes like “G-protein beta subunit binding”, “G-protein Alpha Subunit Binding”, “Gtpase Activating Protein Binding”, and “Morphogenesis” were associated with proteins downregulated in RGS14 KO (Fig. 3D, left), which is unsurprising as we and others have shown the capacity of RGS14 to tightly regulate GPCR and G-protein activation (Harbin et al., 2021) and dendritic spine morphology (Evans et al., 2018b). However, we were surprised to find GO terms like “Response to Prostaglandin Stimulus”, “Nitric-Oxide Synthase Regulator Activity”, and “Intrinsic Pathway for Apoptosis” to be associated with downregulated proteins in RGS14 KO hippocampi (Fig 3D, left), suggesting RGS14 may influence nitric oxide levels and regulate prostaglandin and cellular stress signals. Proteins that were upregulated in RGS14 KO hippocampi after SAL were associated with lipid membrane dynamics and membrane transport (e.g. “Regulation of Membrane Lipid Distribution”, “ATPase-Coupled Lipid Transport”, “Transport Across Blood-Brain Barrier”), receptor tyrosine signaling (e.g. “Receptor Tyrosine Phosphatases”, “EGFR Downregulation”), and acetylcholine neurotransmission (“ACh neurotransmitter Release”) (Fig. 3D, right). We and others have demonstrated RGS14 interacts with small monomeric G proteins like H-Ras to modulate MAPK/ERK signaling (Shu et al., 2010; Willard et al., 2009), making it plausible that RGS14 KO would alter tyrosine receptor signaling dynamics within CA2 pyramidal cells. Although RGS14 has never been demonstrated to modulate lipid dynamics or distribution, blood brain barrier transport, or acetylcholine release, this data demonstrates loss of RGS14 may act upstream to dysregulate a number of cellular pathways that contribute to KA seizure susceptibility in RGS14 KO mice. However, it should be noted that only 8 upregulated proteins were used in this GO analysis, so it may be unlikely that RGS14 majorly affects all of these cellular processes.

To determine alterations in cellular processes caused by loss of RGS14 induction following seizure activity, we performed GO analysis 1 day after KA-SE. After KA treatment, significantly downregulated proteins in RGS14 KO hippocampi are associated with GO terms “Diol Biosynthetic Process”, “Diol Metabolic Process”, “Aldehyde Catabolic Processes”, “Oxidoreductase Activity on Ch-Ch Group”, and “Fatty Acid Metabolism” (Fig. 3E, left), suggesting possible dysregulation of cellular metabolism in RGS14 KO mice following seizure activity. Strikingly, the most enriched GO term associated with proteins downregulated in KO mice was “Eicosanoid Metabolism by COX.” COX-2 is induced after seizure activity and its conversion of arachidonic acid to prostaglandins that contribute to epileptic pathological responses (Rojas et al., 2019). Proteins that are upregulated in RGS14 KO hippocampi were associated with GO terms “Regulation of Intrinsic Apoptosis”, “Positive Regulation of ATP Metabolism”, “Oxygen Binding”, “Antioxidant Activity”, and “Mitochondrial Alpha-Ketoglutarate Dehydrogenase Complex” (Fig. 3E, right). Proteins that localize to the mitochondria or are involved in cellular metabolic regulation are highlighted in the volcano plot (Fig. 3A). Of interest, DEPs in RGS14 KO at baseline include complex I (Ndufaf4), III (Uqcr11), and V (Atp5md, Atp5f1c) components of the electron transport chain (ETC), and DEPs in RGS14 KO following KA-SE include a subunit of the alpha-ketoglutarate dehydrogenase complex Bckdhb and the mitochondrial superoxide dismutase SOD2 (also known as MnSOD). All of these proteins are critical for proper mitochondrial function, and their altered expression in the RGS14 KO hippocampus may exacerbate oxidative stress and seizure pathology (Folbergrova and Kunz, 2012; Shin et al., 2011).

Complimentary to this analysis, we also determined DEPs altered after KA-SE (between treatment) in either WT or RGS14 KO hippocampi (within genotype) (Fig. 4A, B). We found 69 DEPs (34 upregulated, 35 downregulated) in WT mice and 50 DEPs (27 upregulated, 23 downregulated) in RGS14 KO mice, where only 8 proteins were common between the genotypes (Fig. 4C) (complete list of DEPs found in Supplemental Table S2). Again, we used GO analysis on each set of DEPs (Fig. 4D, E), and we compared and contrasted the two GO analyses to provide further insight into KA-induced changes in the RGS14 KO hippocampus relative to WT. Proteins associated with gene expression, protein translation, and protein folding were commonly upregulated in both WT and RGS14 KO (Fig 4D, 4E), which is expected as neuronal activity promotes transcription and translation (Fernandez-Moya et al., 2014; Yap and Greenberg, 2018). In contrast, while upregulated proteins in WT were associated with the GO term “Protein Localization To Mitochondria” (Fig. 4D), upregulated proteins in RGS14 KO lacked this ontology and instead were associated with GO terms “Regulation of Response to Oxidative Stress” and “Cellular Response to Stress”, further implicating that RGS14 may be important in regulating mitochondrial response to seizure activity and loss of RGS14 could promote oxidative stress.

### 3.4. RGS14 localizes to mitochondria in CA2 PCs and reduces mitochondrial respiration in vitro

Among the protein pathways most affected by loss of RGS14, those associated with mitochondrial function and oxidative stress were most surprising and of interest. To explore a possible role for RGS14 in mitochondrial function, we first tested whether RGS14 localizes to the mitochondria in area CA2 PCs. We used immunogold-labeled electron microscopy (EM) to visualize RGS14 in subcellular structures within area CA2 PCs of untreated WT mice (Fig. 5A). Immunogold labeled RGS14 particles are indicated by spherical black staining in proximal dendrites (Fig. 5A, top left and bottom left panels) or cell bodies (Fig. 5A, top center and far right panels). Quite unexpectedly, the electron micrographs depict well-defined immunogold labeling of RGS14 in the external and crista membranes of mitochondria in both dendrites and cell bodies of CA2 pyramidal cells (Fig. 5A). Non-specific immunogold labeling demonstrates a lack of non-specific staining of mitochondria by secondary antibody (Fig. 5B), indicating specific localization of RGS14 in mitochondria of CA2 pyramidal neurons. To determine if RGS14 has an impact on mitochondrial function, we measured oxygen consumption rate (OCR) under basal conditions and following pharmacological inhibition of the ETC in HEK293T cells. We found that cells expressing RGS14 (Fig. 5E) have an overall lower OCR profile compared to cells not expressing RGS14 (Fig. 5C), indicating that RGS14 reduces mitochondrial respiration and is consistent with a previous report that RGS14 had effects on mitochondrial respiration in brown adipose tissue (Vatner et al., 2018). Upon further analysis, we found that RGS14 reduces basal respiration without affecting proton leak, ATP-linked respiration, spare capacity, and non-mitochondrial respiration (Fig. 5D). Together, this data supports a role for RGS14 in the regulation of mitochondrial function, possibly in area CA2 where metabolic function and mitochondrial gene expression is enriched (Farris et al., 2019).

**Figure 5.**
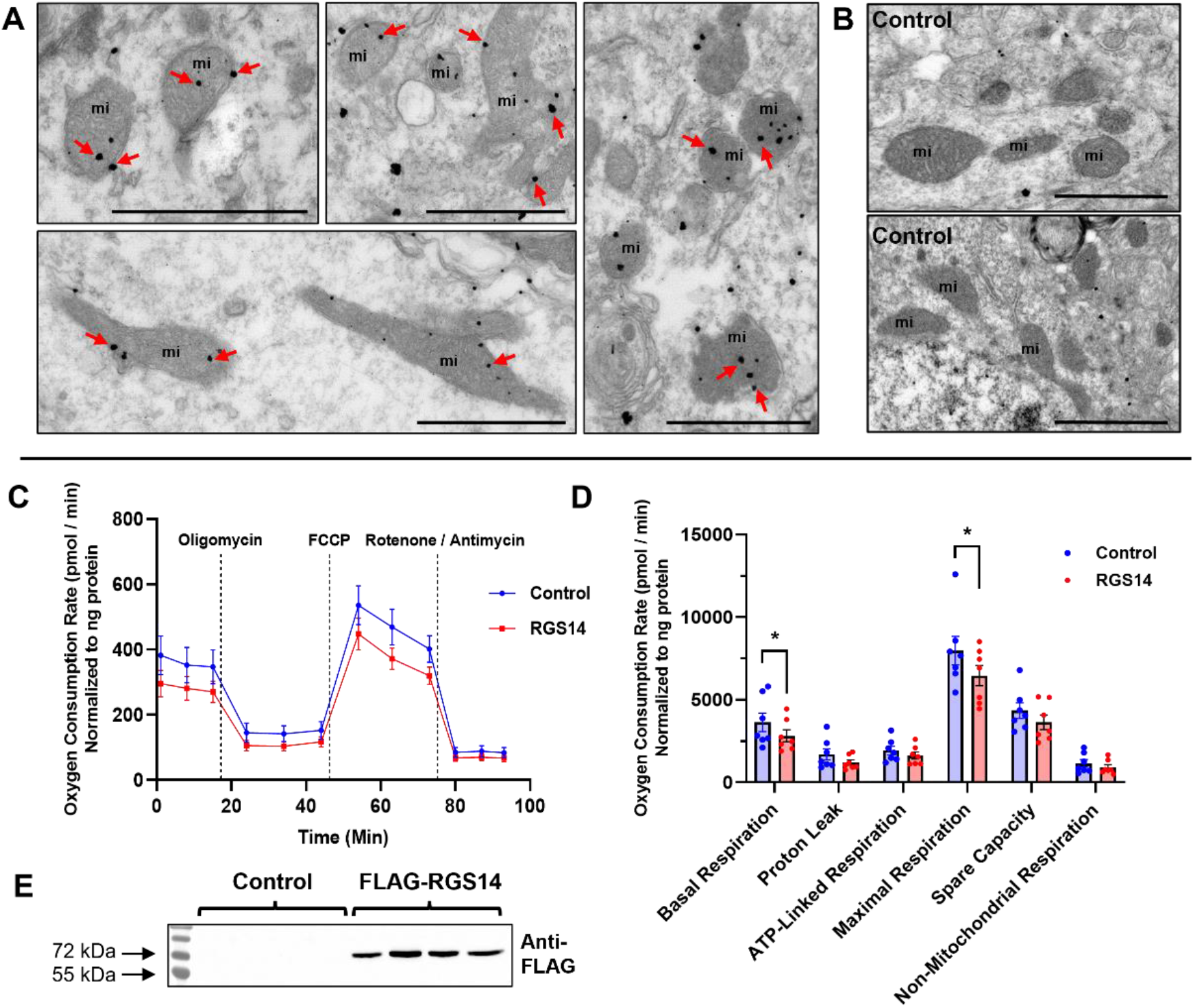
RGS14 localizes to mitochondria in CA2 PCs and reduces mitochondrial respiration *in vitro*. (A) Electron micrographs of CA2 hippocampal region showing RGS14 immunogold labeling in mitochondria. *Top left* and *bottom left* panels depict labeling in dendritic processes, while *center* and *right* panels show neuronal cell body labeling. Gold labeling can be found both on the external and crista membranes of the mitochondria (red arrows). (B) Negative control micrographs showing non-specific immunogold background labeling from sections incubated without the RGS14 antibody. Note the lack of labeling over mitochondria. Scale bars: 1 μm. (C) Normalized oxygen consumption rate plotted over time in HEK293T cells expressing pcDNA3.1 (control) or FLAG-RGS14 (RGS14) during a mitochondrial stress test. Measurements start at baseline and are followed by sequential treatments of cells with mitochondrial inhibitors oligomycin, FCCP, and rotenone/antimycin A. (D) Quantification of parameters derived from panel (C) that are representative mitochondrial function during the assay (n = 7 per transfection condition). (E) Representative Western blot verifying overexpression of RGS14 in HEK293T cells transfected with FLAG-RGS14 compared to pcDNA3.1 (control). *Statistical Analysis:* (D) Unpaired t-tests to compare mean OCR between control vs RGS14 groups (basal respiration, 3627.11 ± 297.19 vs 2819.16 ± 196.95, *p < 0.05; maximal respiration, 7965.98 ± 430.71 vs 6458.40 ± 324.92, *p < 0.05). Error bars represent the SEM. FCCP, carbonilcyanide p-trifluoromethoxyphenylhydrazone. (1.5 column)

### 3.5. RGS14 is necessary for SOD2 induction in area CA2 and prevents 3-nitrotyrosine accumulation

Superoxide and other reactive metabolites are generated by seizure activity, and this seizure-induced oxidative stress plays an important role in mediating seizure pathology (Puttachary et al., 2015). SOD2 is the primary enzyme that detoxifies mitochondrial superoxide, and its altered expression or activity can be deleterious (Holley et al., 2011). Loss of SOD2 exacerbates KA-induced oxidative stress, hippocampal hyperexcitability, and cell death, whereas SOD2 overexpression ameliorates these effects (Liang et al., 2000; Liang and Patel, 2004). We noted that SOD2 was significantly upregulated in RGS14 KO hippocampi after KA as identified by our proteomics (Fig. 3A and Supplemental Table S1).

Because RGS14 localizes to mitochondria in area CA2 (Fig. 5) and loss of RGS14 alters mitochondrial superoxide metabolic pathways (Figs. 3 and 4), we examined the impact of RGS14 loss and seizure on SOD2 protein expression and oxidative stress in hippocampal area CA2 (Fig. 6). We used IHC to quantify SOD2 expression across the hippocampus one day after saline or KA-SE (Fig. 6A). In area CA2, SOD2 expression in both cell bodies (SP) and proximal dendrites (SR) was noticeably similar between WT (Fig. 6A, top left) and RGS14 KO (Fig. 6A, bottom left) after saline treatment. Contrary to our proteomic results, we observed a remarkable increase of SOD2 expression in WT CA2 (Fig. 6A, top right) that was not evident in RGS14 KO CA2 (Fig. 6A, bottom right). In CA2 cell bodies (SP), SOD2 expression was significantly altered by genotype, treatment, and an interaction between the two (Fig. 6B), where post-hoc analysis revealed higher levels of SOD2 in KA-treated WT mice compared to SAL-treated WT CA2 or KA-treated KO CA2. Although there was an apparent increase in SOD2 in CA2 dendrites (SR) in KA-treated WT mice relative to the other groups, these results were not statistically significant (Fig. 6C). In area CA1, we found SOD2 expression to be opposite of area CA2, where SOD2 expression is increased in RGS14 KO compared to WT (Fig. S2). This data suggests RGS14 could influence superoxide levels in the hippocampus by modulating SOD2 expression and also helps explain the paradoxical results between our proteomics and IHC, as whole hippocampi were used for proteomics and the volume of CA2 is considerably smaller than CA1 (Iglesias et al., 2015).

**Figure 6.**
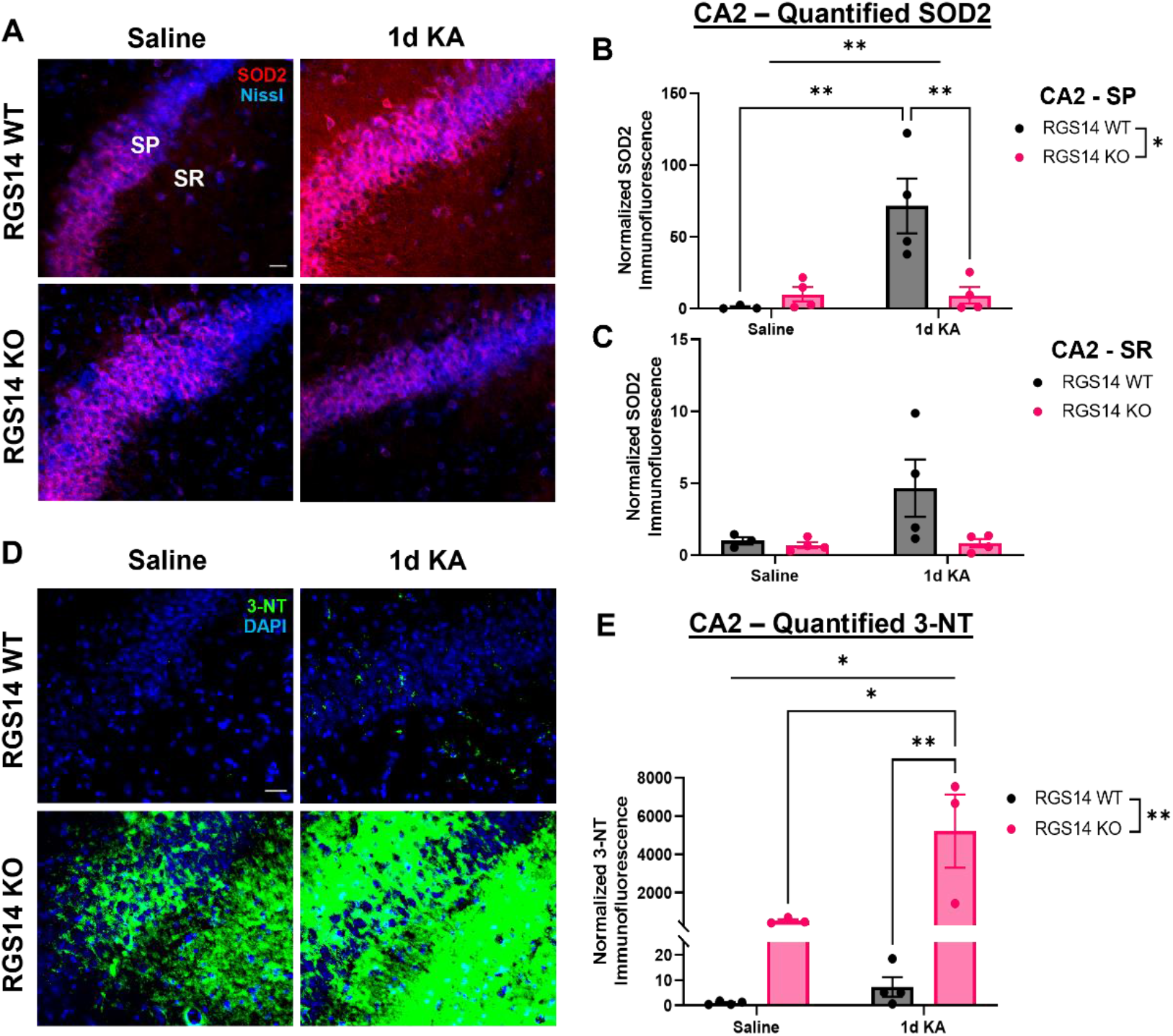
RGS14 is necessary for SOD2 induction in area CA2 and prevents 3-nitrotyrosine accumulation. (A) Representative IHC images of superoxide dismutase 2 (SOD2) expression in WT and RGS14 KO CA2 one day after saline or KA-SE. Note the induction of SOD2 expression after KA-SE in WT but not RGS14 KO CA2. SP and SR represent regions of analysis for cell body (SP) and dendritic (SR) layers of CA2. Scale bar, 20 μm. (B) Mean SOD2 immunofluorescence in the cell bodies (SP) of CA2 expressed relative to saline-treated WT mean (SAL: WT, n=3, 1.00 ± 0.76; RGS14 KO, n=4, 10.09 ± 4.93; 1d KA: WT, n=4, 71.55 ± 19.06; RGS14 KO, n=4, 9.28 ± 5.76). (C) Mean SOD2 immunofluorescence in the proximal dendrites (SR) of CA2 expressed relative to saline-treated WT mean (SAL: WT, 1.00 ± 0.26; RGS14 KO, 0.70 ± 0.21; 1d KA: WT, 4.65 ± 2.00; RGS14 KO, 0.84 ± 0.29). (D) Representative IHC images of 3-nitrotryosine (3-NT) staining in WT and RGS14 KO CA2 one day after saline or KA-SE. 3-NT is detected in striking abundance in RGS14 KO mice compared to WT mice, an effect that is exacerbated following KA-SE. Scale bar, 20 μm. (E) Mean immunofluorescence of 3-NT staining in area CA2 expressed relative to saline-treated WT mean (SAL: WT, n=4, 1.00 ± 0.32; RGS14 KO, n=3, 532.56 ± 83.88; 1d KA: WT, n=4, 7.33 ± 3.86; RGS14 KO, n=3, 5213.93 ± 1908.17). *Statistical analysis:* (B, C, E) Two-way ANOVA with Tukey post-hoc comparisons used to compare group means (B, two-way ANOVA, F = 5.68 for genotype, *p < 0.05; F = 9.77 for treatment, *p < 0.05; F = 10.23 for interaction, **p < 0.01; Tukey’s, WT SAL vs WT KA, **p < 0.01; WT KA vs KO KA, **p < 0.01) (E, two-way ANOVA, F = 12.89 for genotype, **p < 0.01; F = 8.61 for treatment, *p < 0.05; F = 8.56 for interaction, *p < 0.05; Tukey’s, KO SAL vs WT KA, *p < 0.05; WT KA vs KO KA, **p < 0.01). Error bars represent the SEM. SP, stratum pyramidale; SR, stratum radiatum. 3-NT, 3-nitrotyrosine. (1.5 column)

To determine if loss of RGS14 increases superoxide production and exacerbates oxidative stress, we evaluated the expression of hippocampal 3-nitrotyrosine (3-NT) using IHC one day after saline or KA-SE. Tyrosine oxidizes to 3-NT when exposed to peroxynitrite, a highly reactive species that forms from superoxide (O_2_^-^) and nitric oxide (NO). 3-NT is commonly used as a marker for oxidative stress and can greatly impact protein function and signaling (Bandookwala and Sengupta, 2020; Campolo et al., 2020). Quite remarkably, we observed a dramatic increase of 3-NT staining in CA2 of RGS14 KO mice relative to WT after saline or KA-SE. (Fig. 6D). Quantification and comparison of 3-NT staining in area CA2 revealed a significant effect of genotype, treatment, and an interaction between treatment and genotype, suggesting loss of RGS14 and KA-SE influence the abundance of 3-NT in CA2 (Fig. 6E). Although post-hoc analysis showed no difference of CA2 3-NT staining between WT and RGS14 KO after saline treatment, we did find 3-NT abundance in KO CA2 was increased after KA compared to SAL-treated KO CA2 and KA-treated WT CA2) (Fig. 6D). Although RGS14 modulates SOD2 expression in CA1, we observed no genotype or treatment differences in 3-NT staining in this area (data not shown). Overall, these results provide compelling evidence that RGS14 limits excessive amounts of superoxide and prevents accumulation of 3-NT in area CA2 likely by regulating some aspects of mitochondrial function and promoting SOD2 expression after KA.

### 3.6. Loss of RGS14 exacerbates hippocampal neurodegeneration one day following KA-SE

Hippocampal neurodegeneration is a major pathological hallmark of TLE in patients and animal models, including KA-SE (Levesque and Avoli, 2013; Thom, 2014). In KA-SE, neurodegeneration is primarily observed in area CA3 PCs, where seizure activity originates and neuronal damage occurs from hyperexcitation (Hu et al., 1998). Because area CA2 modulates hippocampal excitability and loss of RGS14 increases CA2 PC excitability and sensitizes mice to behavioral seizures by KA (Fig. 1) (Boehringer et al., 2017; Lee et al., 2010), potentially by influencing oxidative stress and mitochondrial function in the area (Figs. 3–6), we hypothesized that RGS14 KO mice would have more severe neurodegeneration in the hippocampus, possibly in CA2 PCs that are typically resistant to such injury (Hatanpaa et al., 2014; Steve et al., 2014). As such, we assessed hippocampal neurodegeneration in dorsal hippocampal sections one day following KA-SE, collected from WT (Fig. 7A) and RGS14 KO (Fig. 7B) mice using Flurojade B (FJB), a sensitive dye that labels degenerating neurons after SE (Schmued et al., 1997). Although we saw no or inconsistent evidence of FJB labeling in areas CA1, CA2, or DG regardless of genotype (data not shown), there was clear and consistent FJB labeling of CA3 pyramidal cells in WT (Fig. 7A, C) and RGS14 KO (Fig. 7B, D) mice. Strikingly, there were considerably more FJB+ CA3 neurons in RGS14 KO mice compared to WT mice. (Fig. 3E). This data supports a neuroprotective role for RGS14 following SE and is consistent with its protective capacity in the heart and liver after insult.

**Figure 7.**
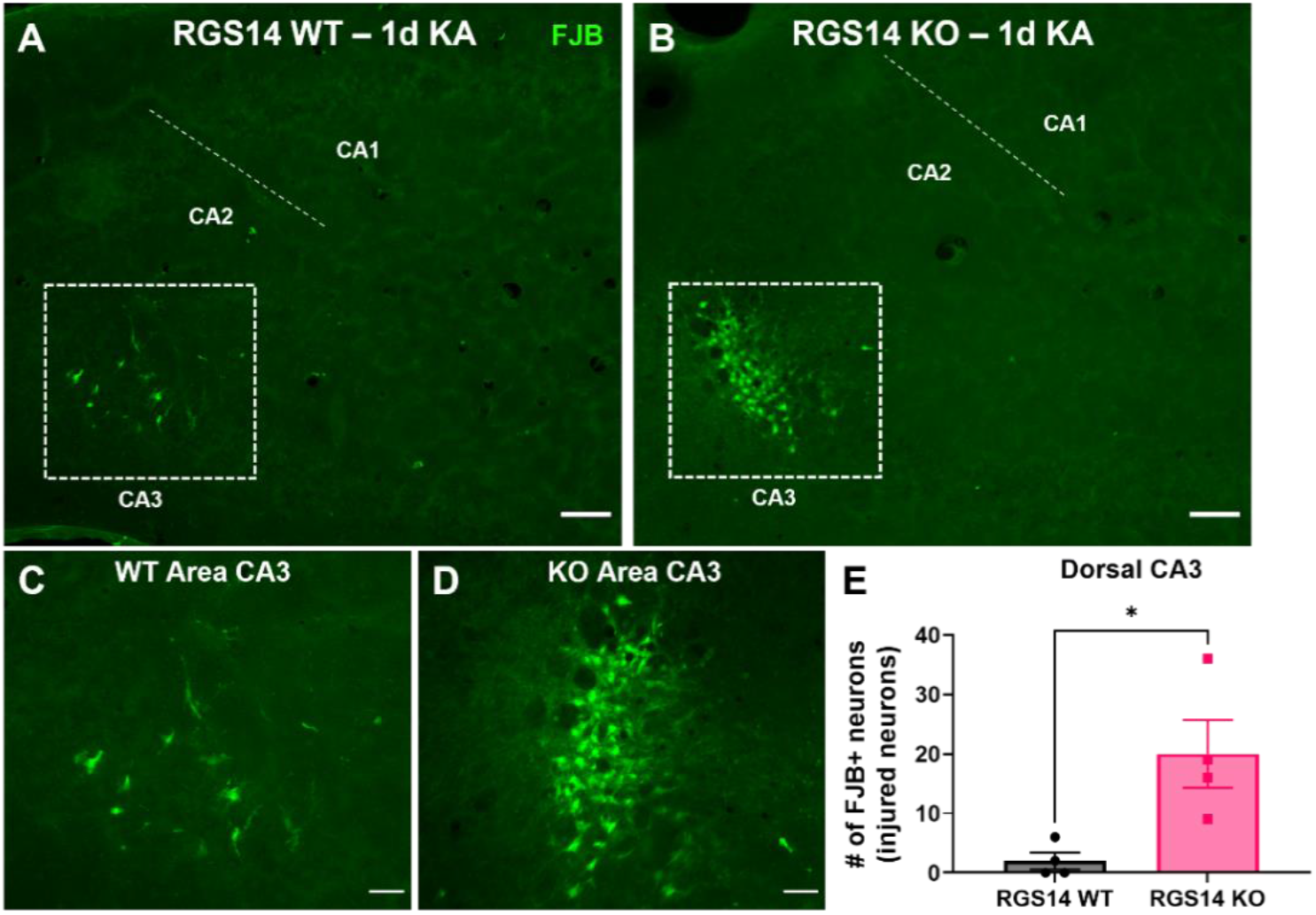
Loss of RGS14 exacerbates hippocampal neuronal injury after KA-SE. (A-B) Representative FluroJade B (FJB) staining in WT (A) and RGS14 KO (B) dorsal hippocampi 1 day following KA-SE (A). Dashed box represents the injured region of area CA3. Scale bar, 50 μm. (C-D) Representative images of area CA3 illustrates increased number of degenerating pyramidal cells in RGS14 KO mice (D) compared to WT mice (C). Scale bar, 20 μm. (E) Mean number of FJB+ neurons in area CA3 of the dorsal hippocampus after KA-SE (n = 4 per genotype) (WT, 2.00 ± 1.41; RGS14 KO, 20.00 ± 5.73). *Statistical Analysis:* (E) Unpaired t-test was used to compare mean number of FJB+ neurons (WT vs RGS14 KO, *p < 0.05). Error bars represent the SEM. (1 column)

### 3.7. RGS14 is required for microglial recruitment and activation one day after KA-SE

Hippocampal microgliosis proceeds SE, where microglia are recruited to sites of hyperexcitability and are activated by a variety of factors that result in morphological change (Hiragi et al., 2018). As a final examination of seizure pathology, we sought to determine if loss of RGS14 altered microglial recruitment and activation to the hippocampus following KA-SE. To test this hypothesis, we performed IHC using IBA1 as a marker to specifically label microglia and quantify microglial number and size in the hippocampus of WT and RGS14 KO mice one day after saline or KA-SE (Fig. 8A). Overall, we observed no visible differences in microglial number or morphology between saline-treated WT and RGS14 KO (Fig. 8A, top panels). However, while there was a noticeable increase in IBA1 immunoreactivity in WT hippocampus after KA-SE, we observed a remarkable lack of increase in IBA1 immunoreactivity in RGS14 KO hippocampus after KA-SE (Fig. 8A, bottom panels), suggesting RGS14 could mediate microglial activation.

**Figure 8.**
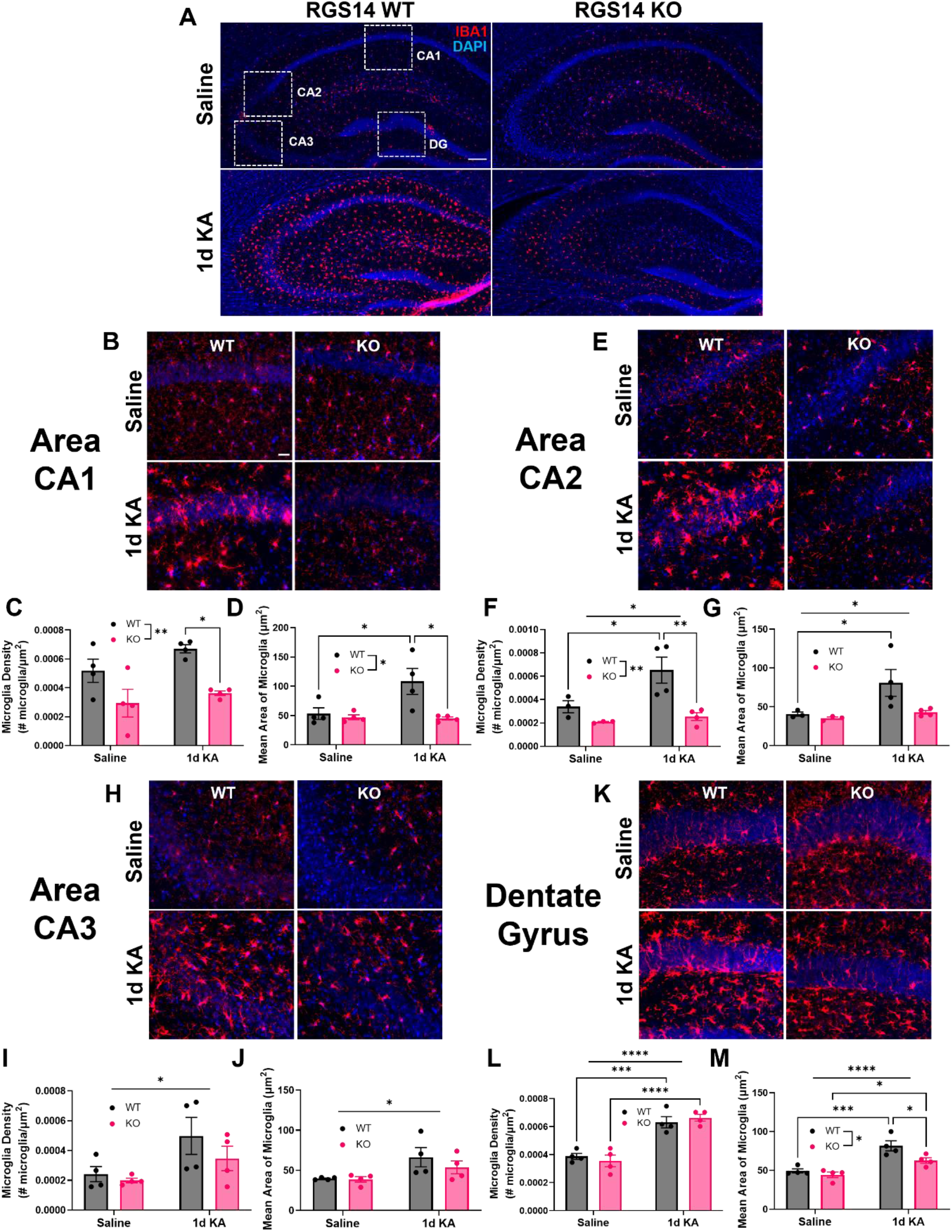
RGS14 is required for microglial recruitment and activation in hippocampal areas CA1 and CA2. (A) Representative IHC images of ionized calcium binding adaptor molecule 1 (IBA1) as a marker of microglia and DAPI (nuclei marker) in the dorsal hippocampus of WT and RGS14 KO mice one day after saline or KA-SE. Scale bar, 100 μm. (B, E, H, K) Representative IHC images of IBA1 expression in area CA1 (B), CA2 (E), CA3 (H), and DG (K) in WT and RGS14 KO mice one day after saline or KA-SE. Scale bar, 20 μm. (C) Mean microglial density for CA1 (# microglia/μm^2^ e-04) (SAL: WT, 5.19 ± 0.80; KO, 2.94 ± 0.96; 1d KA: WT, 6.71 ± 0.28; KO, 3.63 ± 0.16). (D) Mean area (size) of IBA1+ microglia in CA1 (μm^2^) (SAL: WT, 53.19 ± 9.85; KO, 47.09 ± 4.22; 1d KA: WT, 108.31 ± 22.13; KO, 45.02 ± 2.60). (F) Mean microglial density for CA2 (# microglia/μm^2^ e-04) (SAL: WT, n =3, 3.39 ± 0.52; KO, n=3, 2.08 ± 0.05; 1d KA: WT, 6.52 ± 1.11; KO, 2.54 ± 0.33). (G) Mean area (size) of IBA1+ microglia in CA2 (μm^2^) (SAL: WT, n=3, 40.61 ± 2.68; KO, n=3, 35.34 ± 1.61; 1d KA: WT, 80.84 ± 17.26; KO, 42.91 ± 2.37). (I) Mean microglial density for CA3 (# microglia/μm^2^ e-04) (SAL: WT, 2.42 ± 0.51; KO, 1.99 ± 0.15; 1d KA: WT, 4.99 ± 1.24; KO, 3.47 ± 0.82). (J) Mean area (size) of IBA1+ microglia in CA3 (μm^2^) (SAL: WT, 53.19 ± 9.85; KO, 47.09 ± 4.22; 1d KA: WT, 108.31 ± 22.13; KO, 45.02 ± 2.60). (L) Mean microglial density for DG (# microglia/μm^2^ e-04) (SAL: WT, 3.87 ± 0.21; KO, 3.55 ± 0.42; 1d KA: WT, 6.31 ± 0.40; KO, 6.63 ± 0.25). (M) Mean area (size) of IBA1+ microglia in CA1 (μm^2^) (SAL: WT, 49.37 ± 2.61; KO, 44.44 ± 3.68; 1d KA: WT, 81.71 ± 6.50; KO, 62.72 ± 3.60). n=4 per genotype and condition unless noted otherwise. *Statistical analysis:* Two-way ANOVA with Tukey’s or Sidak’s post-hoc comparisons was used to compare mean microglial density (C, F, I, L) and area (D, G, J, M) between treatment and genotype (C, two-way ANOVA, F = 16.95 for genotype, **p < 0.01; Sidak’s, WT KA vs KO KA, *p < 0.05) (D, two-way ANOVA, F = 7.87 for genotype, *p < 0.05; F = 5.35 for interaction, *p < 0.05; Tukey’s, WT SAL vs WT KA, *p < 0.05; WT KA vs KO KA, *p < 0.05) (F, two-way ANOVA, F = 13.41 for genotype, **p < 0.01; F = 6.22 for treatment, *p < 0.05; Tukey’s, WT SAL vs WT KA, *p < 0.05; WT KA vs KO KA, **p < 0.01) (G, two-way ANOVA, F = 5.29 for treatment, *p < 0.05; Sidak’s, WT SAL vs WT KA, *p < 0.05) (I, two-way ANOVA, F = 6.58 for treatment, *p < 0.05) (L, two-way ANOVA, F = 7.81 for treatment, *p < 0.05) (B, two-way ANOVA, F = 68.49 for treatment, ****p < 0.0001; Sidak’s, WT SAL vs WT KA, ***p < 0.001; KO SAL vs KO KA, ****p < 0.0001)) (M, two-way ANOVA, F = 7.58, *p < 0.05; F = 33.93 for treatment, ****p < 0.0001; Tukey’s, WT SAL vs WT KA, ***p < 0.001; KO SAL vs KO KA, *p < 0.05; WT KA vs KO KA, *p < 0.05). Error bars represent the SEM. (2 column)

To determine microglia reactivity (increase in number and size of microglia) and potential subregional differences in microglial activation between WT and RGS14 KO hippocampi, we quantified the density of IBA1+ microglia (number of microglia/area of analysis) and area of IBA1+ microglia for each hippocampal subregion. In CA1 (Fig. 8B), we found a significant effect of genotype on microglia density with significantly more microglia in WT CA1 compared to RGS14 KO CA1 after KA-SE (Fig. 8C). Additionally, there was a significant effect of genotype and interaction between treatment and genotype on microglia size in CA1, where microglia were significantly larger in KA-treated WT mice compared to SAL-treated WT or KA-treated RGS14 KO (Fig. 8D). In CA2 (Fig. 8E), we observed a significant effect of treatment and genotype on microglia density, where there were significantly more microglia in KA-treated WT CA2 compared to SAL-treated WT or KA-treated RGS14 KO CA2 (Fig. 8F). Subsequently, there was an effect of treatment on microglia size in CA2, where only WT microglia increased in size following KA-SE (Fig. 8G). This was a surprising result, as we expected worsened neuronal injury and behavioral seizure response would be associated with increased microgliosis (Heo et al., 2006). Additionally, RGS14 expression in glia has not been demonstrated, which could indicate a neuronal mechanism of promoting microgliosis. However, this suggests that RGS14 is necessary for microglia activation after KA-SE in hippocampal area CA1 and CA2, where RGS14 expression was coincidentally found to be upregulated (Fig. 2).

In contrast to area CA1 and CA2, we found that IBA1+ microglia were similarly activated between WT and RGS14 KO in area CA3 (Fig. 8H). In CA3, there was only a significant effect of treatment on IBA1+ microglia density (Fig. 8I) and average IBA1+ microglia area (Fig. 8J) with no significant diferences between groups in post-hoc analysis. In the DG (Fig. 8K), KA-SE increased microglia density regardless of genotype (Fig. 8L). We also found a significant effect of both treatment and genotype on average IBA1+ microglia area in the DG (Fig. 8M). Although microglia from both WT DG and RGS14 KO DG were significantly larger after KA-SE compared to saline counterparts, WT microglia from DG were slightly but significantly larger than KO DG microglia after KA-SE (Fig. 8M). This data suggests RGS14 is required for microgliosis in area CA1 and CA2 but not CA3 or DG following KA-SE, a novel finding for a protein whose expression is restricted to neurons in the brain, and establishes a clear role for RGS14 in determining early seizure pathology.

## 4. Discussion

Our group and others have shown that RGS14 is integral for area CA2 pyramidal cell (PC) signaling and function (Montanez-Miranda et al., 2022). Based on recent reports of the protective capacity of RGS14 in peripheral tissue (Li et al., 2016; Zhang et al., 2022) and the newly appreciated contribution of area CA2 in TLE (Freiman et al., 2021; Haussler et al., 2016; Kilias et al., 2022; Whitebirch et al., 2022; Wittner et al., 2009), we hypothesized that RGS14 may serve a protective role in area CA2 following seizure insult. Here, we show that loss of RGS14 sensitizes mice to the behavioral effects of KA-SE demonstrated by expedited entry into SE and mortality. In response to seizure activity, RGS14 expression in WT mice is upregulated in the hippocampus, particularly in area CA2 and CA1 PCs. Using proteomics to assess molecular consequences of loss of RGS14, we saw an array of proteins involved with mitochondrial metabolism that differed between WT and RGS14 KO hippocampi following KA-SE. Of note, we observe for the first time that RGS14 localizes to mitochondria of PCs in area CA2 and that add-back of RGS14 to cells reduces mitochondrial respiration *in vitro*. Loss of RGS14 promotes the marked accumulation of 3-nitrotyrosine (3-NT, a marker of oxidative stress) and prevents seizure induction of SOD2 (a mitochondrial enzyme that detoxifies the reactive metabolite superoxide) in CA2 PCs, suggesting exacerbated oxidative stress in the region. Mice lacking RGS14 exhibit increased neuronal injury in area CA3 PCs (but not CA2) while lacking microgliosis in areas CA1 and CA2 following KA-SE. Together, these results provide compelling evidence for the idea that RGS14 is a protective modulator of behavioral and pathological seizure response to KA in the hippocampus, likely by influencing the cellular excitability and metabolic profile of CA2 PCs.

### 4.1 Loss of RGS14 primes the hippocampus to excitability and pathology prior to seizure induction

Faster onset of SE and mortality in RGS14 KO suggests enhanced hippocampal excitability and/or propagation of seizure activity from the hippocampus (Fig. 1). CA2 PCs from RGS14 KO mice are intrinsically more excitable than those from WT mice, and, in contrast to WT CA2 PCs that lack CA3-CA2 plasticity (Zhao et al., 2007), CA2 PCs from RGS14 KO express robust synaptic plasticity at these synapses (Lee et al., 2010), which may enhance excitatory drive onto CA2 in RGS14 KO mice. Because epileptiform activity originates in area CA3 following KA injection (Ben-Ari and Cossart, 2000), lost gating of CA3-CA2 synapses and increased CA2 excitability could propagate seizure activity to CA2 and its targets faster in RGS14 KO mice, resulting in faster entry into SE activity and mortality. Because CA2 has a wide, bilateral ouput along and out of the hippocampus (Cui et al., 2013; Mercer et al., 2007), increased excitation onto and out of CA2 in RGS14 KO mice may initiate seizure activity in these regions more rapidly or increase the likelihood of seizure activity, similar to what has been shown in epileptic CA2 (Whitebirch et al., 2022).

Against this interpretation, chronic inhibition of CA2 results in CA3 hyperactivity, epileptiform discharges in CA3, and increased sensitivity to KA-induced behavioral seizures (Boehringer et al., 2017). In this context, CA2 hyperactivity would decrease CA3 activity and epileptiform activity caused by KA in RGS14 KO mice, contrary to our findings here. This discrepancy may reflect RGS14 expression and regulation in other seizurogenic areas like the amygdala, entorhinal, and piriform cortex (Evans et al., 2014; Squires et al., 2021). Alternatively, RGS14 may be more important for controlling CA2 output to propagate epileptiform activity rather than the generation of epileptiform activity within the CA3-CA2 network. Expedited entry into SE and mortality in RGS14 KO mice may reflect quicker seizure propagation from CA2 to other limbic structures that result in convulsive limbic seizures (i.e. Stage 3/SE) and to hindbrain structures that regulate autonomic function, which may disrupt breathing, cardiac function, and bring about death quicker than in WT mice (Ben-Ari and Cossart, 2000; Massey et al., 2014). Therefore, future studies could employ region-specific RGS14 deletion and EEG recording to assess how RGS14 affects epileptiform activity within the hippocampus and seizure propagation to connected structures.

Importantly, the capacity of RGS14 to modulate CA2 activity may be clinically relevant as CA2 has been hypothesized to promote epileptogenesis and generate seizure activity in the absence of CA1 and CA3 (Kilias et al., 2022; Whitebirch et al., 2022), which are susceptible to degeneration in TLE relative to the injury resistant CA2 (Steve et al., 2014). CA2 epileptiform activity has been reported in KA and pilocarpine models of TLE (Kilias et al., 2022; Whitebirch et al., 2022) as well as resected tissue from TLE patients (Wittner et al., 2009). A recent report also demonstrated increased excitability of CA2 after TLE development in mice, where acute inhibition of CA2 reduced seizure frequency in mice that developed recurrent seizures (Whitebirch et al., 2022). This evidence supports the idea that CA2 is involved in promoting seizure activity in TLE, and supports our findings that RGS14 in CA2 PCs would influence excitability of this region and in the hippocampus to modulate behavioral seizures.

### 4.2. Altered protein profiles that influence seizure susceptibility after loss of RGS14

Proteomic analysis of RGS14 KO hippocampus (Fig. 3, 4, S1-2) showed that downregulated proteins were associated with G protein dependent signaling in RGS14 KO (e.g. Rap2c, Gnas), which could reflect aberrant G protein signaling within the hippocampus, and morphogenesis, which was expected as we have shown RGS14 inhibits glutamate-dependent increases in CA2 dendritic spine volume (Evans et al., 2018b). Additionally, the presence of receptor tyrosine-related GO pathways in upregulated proteins of RGS14 KO made sense in the context of previous findings, where RGS14 inhibits receptor tyrosine signaling by binding to MAPK/ERK components like H-Ras (Shu et al., 2010; Vellano et al., 2013). Interestingly, the GO pathway “Response to Prostaglandin Stimulus” was enriched in downregulated proteins caused by RGS14 KO. Prostaglandins exert their effects through a number of GPCRs, including those coupled to Gi/o (Hata and Breyer, 2004), where RGS14 could limit their activity by acting through its GAP/RGS function. Prostaglandins like PGE2 are known to regulate membrane excitability, seizure response, and inflammation in the hippocampus (Chen and Bazan, 2005; Jiang et al., 2013) (Jiang et al., 2013; Rojas et al., 2014), which is coincident with the phenotype of RGS14 KO and may suggest RGS14 (albeit indirectly) control over prostaglandin stimulation. Overall, this data suggests RGS14 limits intracellular signaling, likely in CA2 PCs where it is highly expressed, to decrease CA2 excitability and prevent exacerbated seizure response to KA. Even so, this interpretation is limited by the fact that changes in protein expression detected by proteomics only partially reflect those in CA2 PCs since data sets were obtained from whole hippocampi and global deletion of RGS14 was not limited to CA2. Although findings may reflect broader changes in signaling in the hippocampus due to loss of RGS14, future studies examining cell-type specific deletion and analysis will be needed to more accurately assess RGS14 roles in CA2 to the observed phenotype.

### 4.3. Regulation of mitochondrial function and oxidative stress by RGS14

The most unexpected and intriguing findings of our report is that RGS14 is involved with aspects of metabolism, mitochondrial function, and oxidative stress in the hippocampus (Figs. 3–6). We found RGS14 localizes to mitochondria in CA2 PCs and reduces basal and maximal mitochondrial respiration *in vitro*, which corroborates a previous finding that RGS14 KO increases mitochondrial respiration in brown adipose tissue (Vatner et al., 2018). How RGS14 alters mitochondria function is unknown. RGS14 actions there could reflect upstream regulation of GPCR, MAPK, or calcium signaling at the plasma membrane (Harbin et al., 2021), or at the mitochondria to limit GPCR activation (Benard et al., 2012; Hollinger et al., 2001) or buffer calcium as it does at the membrane (Duchen, 2000; Evans et al., 2018b). How RGS14 localizes to the mitochondria is also uncertain. At least one predictive algorithm for mitochondrial targeting sequences (MTS) suggest that RGS14 may contain an N-terminal MTS (Fukasawa et al., 2015). Alternatively, RGS14 may bind to an effector or chaperone to target the organelle (Schmidt et al., 2010). Future studies will explore these possibilities.

Even though our findings do not directly measure RGS14 effects on mitochondrial respiration in neurons, we observe robust RGS14 localization to mitochondria in CA2 neurons and propose that RGS14 could regulate mitochondrial function in neurons as it does in brown adipocytes (Vatner et al., 2018). Both neurons and brown adipocytes are high in mitochondrial content and must meet intense energy demands for proper organ function (Attwell and Laughlin, 2001; Magistretti and Allaman, 2015; Shinde et al., 2021). Moreso, CA2 PCs exhibit higher metabolic activity, more mitochondrial DNA and mitochondrial-related mRNA transcripts, and plasticity-dependent mitochondrial calcium uptake relative to neighboring CA1 and CA3 (Farris et al., 2019). The fact that RGS14 is highly enriched in CA2 neurons, localizes to mitochondria, and reduces mitochondrial respiration, illustrates that RGS14 is ideally positioned to influence mitochondrial function in CA2.

Tyrosine nitrosylation (3-NT) (elevated in RGS14 KO CA2 (Fig. 6)) is caused by the highly reactive peroxynitrite, which is generated by nitric oxide (NO) and superoxide (O_2_^-^) (Ferrer-Sueta et al., 2018), and suggests enhanced oxidatives stress (Bandookwala and Sengupta, 2020). This finding was supported by our proteomic data where we observed altered expression of proteins involved with the mitochondrial electron transport chain and metabolic processes, apoptosis, and regulation of NO generation and oxidative stress in RGS14 KO hippocampus (Figs. 3, 4). By reducing mitochondrial respiration (Fig. 6), RGS14 may limit the production of O_2_^-^ and subsequent oxidative stress (Turrens, 2003). Additionally, RGS14 suppresses NMDAR-dependent calcium influx, signaling, and plasticity in CA2 dendritic spines (Evans et al., 2018b). Notably, calcium influx leads to mitochondrial O_2_^-^ formation in hippocampal neurons (Hongpaisan et al., 2004), while NMDAR agonism generates NO and O_2_^-^ via activation of NO synthase (NOS) and NADPH oxidase (NOX), respectively, at the membrane (Di Meo et al., 2016). By limiting signaling at the plasma membrane and/or in the mitochondria, there is potential for RGS14 to inhibit NO and/or O_2_^-^ production. Follow up studies will determine if RGS14 modulates O_2_^-^ and/or NO in hippocampal neurons (at baseline or following seizure activity) and utilize pharmacological inhibition to elucidate their source(s).

Furthermore, loss of RGS14 resulted in a marked suppression of SOD2 (the primary mitochondrial O_2_^-^ detoxifying enzyme) in CA2 after KA-SE, which could explain, at least in part, the elevation of 3-NT caused by seizure activity. 3-NT impacts the structure and activity of proteins susceptible to tyrosine nitrosylation, especially those in the mitochondria which can have deleterious effects on its function (Ferrer-Sueta et al., 2018; Radi et al., 2002). Indeed, SOD2 is inactivated by 3-NT modification (Yamakura et al., 1998), which may contribute to further O_2_^-^ production and oxidative stress (Holley et al., 2011). Related, SOD2 deficient mice develop spontaneous seizures and are more sensitive to KA-induced seizures (Liang and Patel, 2004; Liang et al., 2012). In addition to 3-NT effects on mitochondrial enzymes, NO and O_2_^-^ modify the function of ion channels at the membrane that may influence excitability in RGS14 KO CA2 PCs (Bogeski and Niemeyer, 2014; Radi et al., 2002; Spiers and Steinert, 2021). Therefore, loss of SOD2 and elevated 3-NT in area CA2 of RGS14 KO mice likely reflect a cellular environment with elevated levels of O_2_^-^ and/or NO, which would have a major role in promoting oxidative stress, altering cellular excitability, and contributing to seizure generation in CA2.

### 4.4. Possible protective roles of RGS14 in CA2 hippocampus

We found RGS14 protein is upregulated markedly in CA2 and CA1 PCs following seizure (Fig. 2), similar to RGS14 upregulation in hepatocytes that is protective after ischemic-reperfusion injury (Zhang et al., 2022). In the hippocampus, RGS14 induction may serve as negative feedback to limit excessive calcium and aberrant signaling caused by seizure (Evans et al., 2018b; Harbin et al., 2021). Complimentary to its influence on signaling, RGS14 induction likely limits seizure-induced oxidative stress and mitochondrial dysfunction in CA2 PCs. While RGS14 was also upregulated in CA1, we did not see 3-NT accumulation and observed the opposite effect on SOD2 which was increased in CA1 compared to WT. This may reflect compensatory mechanisms in CA1 that properly limit O_2_^-^ generation, 3-NT modification, and oxidative stress despite lacking RGS14 induction. Indeed, the translational and transcriptional profiles in areas CA2 and CA1 differ considerably, suggesting different mechanisms (Farris et al., 2019; Gerber et al., 2019). Therefore, RGS14 likely modulates mitochondrial function and oxidative stress only in CA2 as a mode of protection. Mechanism of RGS14 induction may also underly how protection is afforded. Our unpublished findings show no change in the occupancy of the promoter region or regions upstream of the *RGS14* gene, suggesting RGS14 induction involves translation of RGS14 mRNA rather than transcription. Indeed, a recent report found RGS14 mRNA is highly abundant in the dendrites and cell bodies of CA2 and CA1 PCs, suggesting local translation of mRNA as a mechanism of RGS14 induction (Farris et al., 2019). Future experiments will employ transcriptional and translational inhibition to define the mechanism of RGS14 induction.

As direct evidence of RGS14 neuroprotection, we observed enhanced seizure-induced neuronal injury in CA3 PCs after loss of RGS14 (Fig. 7), which is associated with excessive excitation of CA3 in KA-SE (Ben-Ari and Cossart, 2000; Rusina et al., 2021). Contrary to our hypothesis, we observed no injury in CA2, suggesting RGS14 is dispensable for CA2 resistance likely due to compensatory protection from a number of CA2-specific mechanisms (e.g. strong inhibitory drive, high expression of calcium buffering proteins) (Dudek et al., 2016). The metabolic changes in CA2 due to loss of RGS14 (enhanced 3-NT, decreased SOD2) were not evident in CA3 and RGS14 was not expressed in CA3, suggesting RGS14 modulates CA3 injury via its regulation in CA2. Therefore, we propose that the enhanced neuronal injury in CA3 is due to due to lack of signaling and metabolic regulation in CA2 PCs by RGS14 that may result in a net increase of excitation onto CA3 PCs. As proposed earlier, future studies will need to evaluate RGS14 KO hippocampal excitability during seizure activity to properly determine the mechanism of CA3 injury.

### 4.5. Altered microgliosis and inflammatory pathways following loss of RGS14

Although lasting neuroinflammation is typically associated with worse seizure pathology and outcomes, several studies suggest early microgliosis following seizure is protective in the hippocampus (Eyo et al., 2014; Liu et al., 2020; Wu et al., 2020). Loss of early microglial activation in RGS14 KO is consistent with a neuroprotective role for RGS14. Glial activation by seizure activity is a complex process, including release of factors from neurons, astrocytes, and microglia (Eyo et al., 2017; Patel et al., 2019). Our proteomics indicate an alteration of cyclooxygenase (COX) and prostaglandin signaling in RGS14 KO mice (Fig. 3). Because prostaglandins (e.g. PGE2) and their synthesis by COX-2 are mediators of microgliosis after seizure (Rojas et al., 2019), dysfunction in this pathway may be relevant to the lack of microgliosis in RGS14 KO hippocampus. Additionally, NMDAR-dependent release of ATP helps attract microglia to the site of excitation (Eyo et al., 2014), and RGS14 regulates downstream effects of NMDAR activation (Evans et al., 2018b). Lack of microgliosis in CA1/CA2 in RGS14 KO mice correlates with RGS14 upregulation in WT CA1/CA2, suggesting RGS14 induction is necessary for regional microglia activation, where it could promote release of activating or chemoattractive factors for microglia. Future experiments will seek to unravel the underlying factors (or lack of) that may suppress microgliosis in RGS14 KO hippocampus.

Although some microglial/macrophage populations express RGS14 mRNA (Bennett et al., 2016; Li et al., 2019), we have not observed RGS14 protein expression in microglia or astrocytes in the brain (Evans et al., 2014; Squires et al., 2018) or RGS14 colocalization with IBA1 in CA1 or CA2 at baseline or following KA-SE. Therefore, we believe it is more likely that RGS14 promotes microglial activation via a neuron-dependent mechanism. However, we cannot rule out a priming effect towards astrocytes and/or microglia, where lack of RGS14 in CA2 and CA1 neurons may affect astrocytic or microglial function prior to injury. Thus, deficiency in microglial or astrocytic signaling and function may contribute to the lack of their activation in RGS14 KO (Eyo et al., 2017; Patel et al., 2019). To delineate the neuronal vs glial contributions in the RGS14 KO phenotype, future studies could examine *ex vivo* activation of microglia and astrocytes isolated from WT and RGS14 KO hippocampi.

### 4.6. Conclusions

Overall, our findings suggest that RGS14 is neuroprotective in the hippocampus by dampening behavioral seizures and regulating CA2 PC function by engaging mitochondria oxidative stress pathways as illustrated in our proposed working model (Fig. 9). Consistent with this idea, RGS14 protein is upregulated in area CA2 after KA-induced SE. Mice lacking RGS14 have faster onset of SE and mortality, striking alterations in metabolic and mitochondrial protein expression, enhanced oxidative stress, increased neuronal injury, and region-dependent suppression of microglial activation. Although mechanistic insights into RGS14 roles in seizure response are limited in our study, we show that RGS14 localizes to mitochondria in CA2 hippocampus and regulates mitochondrial respiration as one possible and newly appreciated mechanism for RGS14 protective actions. Additionally, our observed changes in protein sets/pathways provide compelling evidence of altered responses in RGS14 KO mice that are ripe for future study. Unclear is whether these early pathological alterations contribute to the development of spontaneous seizures and mossy fiber sprouting, which are fundamental consequences of TLE, and awaits further study. Future studies also will explore how RGS14 engages the mitochondria to regulate their function. Together, our findings highlight a protective role of RGS14 in area CA2, seizure development, and related pathology, and sheds insight as to how CA2 participates in the development of TLE.

**Figure 9.**
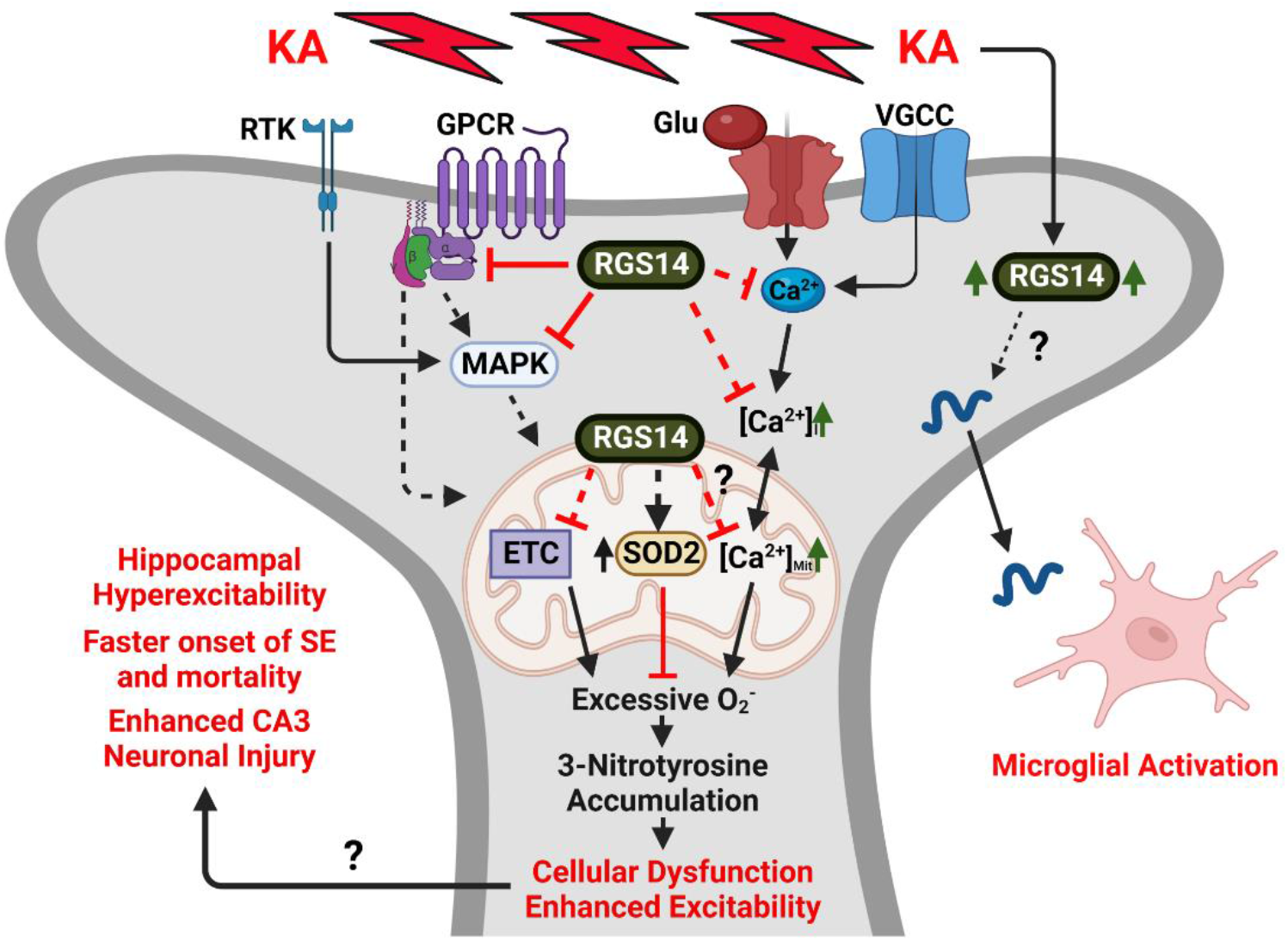
Proposed working model for RGS14 regulation of CA2 pyramidal neurons following seizure. Diagram of a CA2 dendritic spine illustrating known and speculative roles for RGS14 in intracellular regulation. RGS14 at the plasma membrane and in the cytosol can negatively regulate both GPCR- and MAPK-dependent signaling, while buffering calcium transients from NMDA receptors and voltage-gated calcium channels (VGCC) through an unknown mechanism. In addition to membrane and cytosolic spaces, RGS14 localizes to mitochondria in CA2 pyramidal cells. By its effect on signaling and calcium buffering at the membrane/cytosol, its localization to the mitochondria, or through an unknown mechanism, RGS14 is capable of influencing metabolic/mitochondrial protein expression, including SOD2. Although speculative, RGS14 may buffer mitochondrial calcium in a similar fashion to buffering membrane calcium transients of dendritic spines. Regulatory capacity by RGS14 culminates to provide balance of reactive metabolites (O_2_^-^) and mitochondrial function. Additionally, KA seizure activity upregulates RGS14, which may be necessary for promoting microglial activation possibly by the synthesis and release of activating factors. When RGS14 is absent, this leads to overwhelming abundance of 3-nitrotyrosine (3-NT), an indicator of oxidative stress, which may be enhanced after seizure due to the loss of SOD2 upregulation and 3-NT inactivation of SOD2. Exacerbated oxidative stress in RGS14 KO likely causes cellular dysfunction and alters the excitatory properties of CA2 pyramidal cells, leading to the enhanced hippocampal excitability, accelerated onset of SE and mortality, and worse neuronal injury in CA3. GPCR, G-protein coupled receptor; RTK, receptor tyrosine kinase; MAPK, mitogen-activated protein kinase; NMDA, N-methyl-D-aspartate; KA, kainic acid; Glu, glutamate; VGCC, voltage-gated calcium channel; Ca^2+^, calcium; SOD2, superoxide dismutase 2; O_2_^−^, superoxide. Created with BioRender.com. (1.5 column)

## Acknolwedgements

The authors would like to acknowledge Asheebo Rojas for being a valuable intellectual resource during development and execution of experiments, Nicholas Varvel for assistance with behavioral seizure scoring, and Thomas Shiu for preliminary seizure induction studies. This work was supported by the National Institutes of Health [R01NS037112 (to J. R. H.), R01GM140632-A1 (to J. R. H and Peter A. Freidman), F31AA029938 (to K. M. C.), R01AA026086 (to S. M. Y), T32GM008602 (to Randy A. Hall)]; Emory Integrated Proteomics Core (EIPC), which is subsidized by the Emory University School of Medicine and is one of the Emory Integrated Core Facilities; and the Georgia Clinical & Translational Science Alliance of the National Institutes of Health [UL1TR002378]. The content is solely the responsibility of the authors and does not necessarily reflect the official views of the National Institutes of Health, the Department of Veterans Affairs, or the US Government.

