## Supplemental Material for "RGS14 is neuroprotective against seizure-induced mitochondrial oxidative stress and pathology in hippocampus"

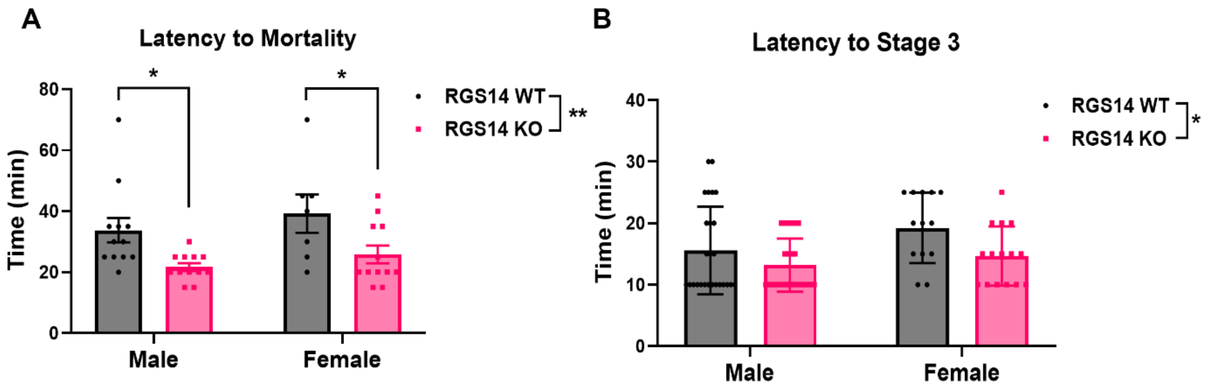


**Supplemental Figure S1. Behavioral sensitivity to KA-SE in RGS14 KO mice is not affected by sex.** (A) Mean latency to mortality in male and female WT and RGS14 KO mice (males: WT, 33.75 ± 4.00 min; RGS14 KO, 21.67 ± 1.28 min) (females: WT, 39.29 ± 6.31 min; RGS14 KO, 25.83 ± 2.94 min). (B) Mean latency to Stage 3 behavioral seizure activity in male and female WT and RGS14 KO mice. (A, B) Two-ANOVA with Sidak post-hoc comparisons was used to compare mean latency to mortality (A; two-way ANOVA, $F = 12.60$ for genotype, $^{**}p < 0.01$; Sidak, WT vs RGS14 KO, $^{*}p < 0.05$) and Stage 3 seizure activity (two-way ANOVA, $F = 6.71$ for genotype, $^{*}p<0.05$). Error bars represent the SEM.

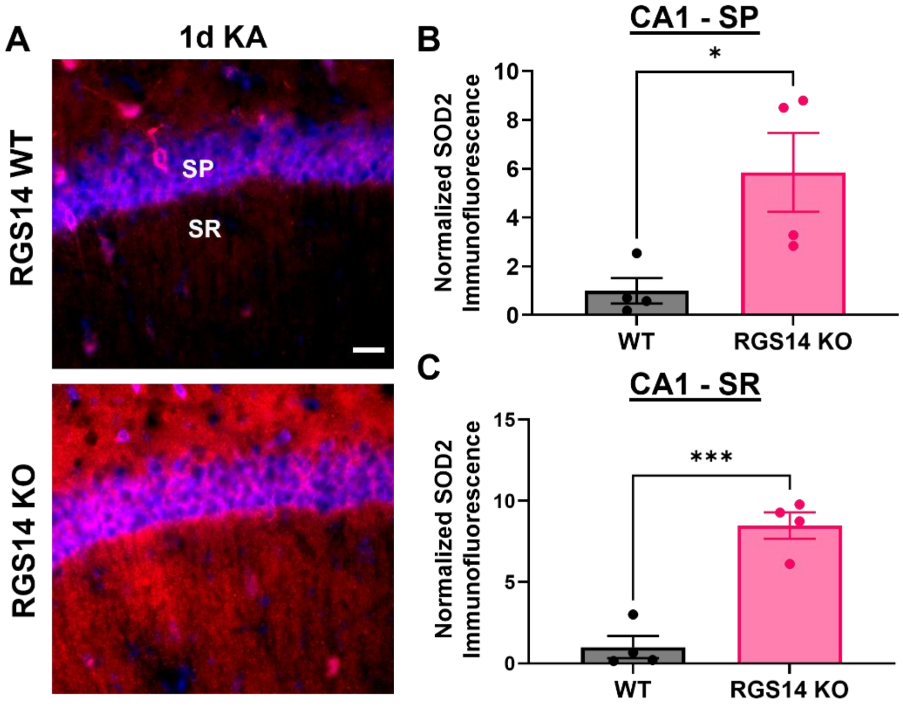


**Supplemental Figure S2. SOD2 is upregulated in area CA1 of RGS14 KO mice one day after KA-SE.** (A) Representative IHC images of superoxide dismutase 2 (SOD2) expression in WT and RGS14 KO CA2 one day after KA-SE. SP and SR represent regions of analysis for cell body (SP) and dendritic (SR) layers of CA2. (B) Mean SOD2 immunofluorescence in the SP layer of CA1 expressed relative to WT mean (WT, 1.00 ± 0.52; RGS14 KO, 5.85 ± 1.62). (C) Mean SOD2 immunofluorescence in the SR layer of CA1 expressed relative to WT mean (WT, 1.00 ± 0.68; RGS14 KO, 8.48 ± 0.81). (B, C) Group means compared using unpaired t-tests (B, WT vs RGS14 KO, *$p < 0.05$) (C, WT, vs RGS14 KO, ***$p < 0.001$). Error bars represent the SEM. SP, stratum pyramidale; SR, stratum radiatum.

**A** **SAL**

Upregulated in KO

| Protein | Adj. p-value | Log2 FC (KO/WT) |
|---|---|---|
| Ehbp1 | 0.00342 | 1.848 |
| Ppfia4 | 0.00146 | 0.707 |
| Them6 | 0.01911 | 0.537 |
| Ocrl | 0.01824 | 0.518 |
| Abcb1a | 0.00534 | 0.381 |
| Ndufaf4 | 0.03588 | 0.309 |
| Sh3kbp1 | 0.02090 | 0.290 |
| Atp5f1c | 0.03784 | 0.167 |

Downregulated in KO

| Protein | Adj. p-value | Log2 FC (KO/WT) |
|---|---|---|
| Rgs14 | 0.00003 | -4.239 |
| Tpm1 | 0.04297 | -1.405 |
| Spag7 | 0.00010 | -1.054 |
| Gfer | 0.00540 | -0.907 |
| Uqcr11 | 0.01041 | -0.899 |
| Rap2c | 0.00878 | -0.808 |
| Nfix | 0.02050 | -0.779 |
| Pigu | 0.03585 | -0.626 |
| Fto | 0.01307 | -0.597 |
| Syt11 | 0.04393 | -0.591 |
| Naa15 | 0.01176 | -0.579 |
| Trappc8 | 0.02520 | -0.537 |
| Naaa | 0.02662 | -0.521 |
| Ggact | 0.04601 | -0.517 |
| Psap | 0.03392 | -0.501 |
| Srrm2 | 0.03583 | -0.496 |
| Ndrg4 | 0.04098 | -0.483 |
| Rab9a | 0.03169 | -0.472 |
| Atp5md | 0.04234 | -0.448 |
| Tiprl | 0.00271 | -0.420 |
| Ddost | 0.01995 | -0.418 |
| Scg5 | 0.03750 | -0.401 |
| Edc4 | 0.03111 | -0.395 |
| Erlin2 | 0.00245 | -0.323 |
| Akt1 | 0.01005 | -0.322 |
| Agfg2 | 0.01151 | -0.319 |
| Scn1b | 0.04604 | -0.310 |
| Dynll1 | 0.04879 | -0.277 |
| Alcam | 0.04788 | -0.263 |
| Ggt7 | 0.04351 | -0.252 |
| G6pdx | 0.02414 | -0.245 |
| Slc4a4 | 0.04152 | -0.190 |
| Ndufv1 | 0.03694 | -0.151 |
| Gnas | 0.02256 | -0.139 |
| Cct5 | 0.04734 | -0.127 |
| Actn1 | 0.01044 | -0.097 |
| Fasn | 0.01851 | -0.087 |

**B** **KA**

Upregulated in KO

| Protein | Adj. p-value | Log2 FC (KO/WT) |
|---|---|---|
| Apob | 0.03976 | 1.116 |
| Ehbp1 | 0.01315 | 0.930 |
| Txndc12 | 0.01455 | 0.825 |
| Bckdhb | 0.04040 | 0.796 |
| Mlst8 | 0.01556 | 0.731 |
| Ppfia4 | 0.01394 | 0.538 |
| Arsb | 0.00106 | 0.488 |
| Cygb | 0.03770 | 0.477 |
| Tmed7 | 0.02162 | 0.477 |
| Trappc8 | 0.04369 | 0.452 |
| Dlgap2 | 0.03178 | 0.419 |
| Acat2 | 0.02662 | 0.385 |
| Sod2 | 0.02007 | 0.329 |
| Sort1 | 0.03027 | 0.303 |
| Ssr1 | 0.01621 | 0.293 |
| Eno1 | 0.03296 | 0.090 |

Downregulated in KO

| Protein | Adj. p-value | Log2 FC (KO/WT) |
|---|---|---|
| Rgs14 | 0.00005 | -4.070 |
| Sf3b1 | 0.03869 | -0.904 |
| Gfer | 0.03674 | -0.851 |
| Fto | 0.00107 | -0.808 |
| Hnrnph3 | 0.03684 | -0.690 |
| Srgap2 | 0.04268 | -0.654 |
| Prrc2c | 0.01850 | -0.647 |
| Gabbr1 | 0.04053 | -0.637 |
| Elmo1 | 0.01756 | -0.627 |
| Stx8 | 0.02297 | -0.551 |
| Hagh | 0.00063 | -0.517 |
| C2cd4c | 0.03492 | -0.508 |
| Trappc13 | 0.01821 | -0.494 |
| Anxa3 | 0.03948 | -0.483 |
| Cacna1e | 0.01062 | -0.472 |
| Ptgr2 | 0.01220 | -0.460 |
| Rgs7bp | 0.00155 | -0.459 |
| Ubr4 | 0.01557 | -0.451 |
| Bola2 | 0.04830 | -0.414 |
| Dgke | 0.04097 | -0.398 |
| Esd | 0.03030 | -0.368 |
| Mpp3 | 0.01045 | -0.311 |
| Acaa1a | 0.02432 | -0.306 |
| Asah1 | 0.02940 | -0.303 |
| Npc1 | 0.01759 | -0.297 |
| Psmc6 | 0.01991 | -0.292 |
| Tiprl | 0.04249 | -0.282 |
| Ncam1 | 0.04696 | -0.282 |
| Sh3kbp1 | 0.03674 | -0.265 |
| Hdhd2 | 0.01715 | -0.257 |
| Spr | 0.04502 | -0.158 |
| Actr1b | 0.01800 | -0.153 |
| Fasn | 0.00172 | -0.119 |

**Supplemental Table 1. Comprehensive list of differentially expressed proteins (DEPs) between WT and RGS14 KO after SAL or KA treatment.** (A) List of upregulated (left) or downregulated (right) DEPs in RGS14 KO hippocampi one day after saline treatment, their adjusted p-values, and the fold change of abundance ($log_2$ FC (KO/WT)). (B) List of upregulated (left) or downregulated (right) DEPs in RGS14 KO hippocampi one day after saline treatment, their adjusted p-values, and the fold change of abundance ($log_2$ FC (KO/WT)). FC, fold change.

**A** **WT**

Upregulated after KA

| Protein | Adj. p-value | Log2 FC (KA/Sal) |
|---|---|---|
| Lrrfip1 | 0.04855 | 2.039 |
| Hspb1 | 0.01937 | 1.367 |
| Dnajb5 | 0.00046 | 1.217 |
| Ybx1 | 0.03460 | 1.132 |
| Vgf | 0.00210 | 0.952 |
| Castor2 | 0.04962 | 0.932 |
| Lsm3 | 0.03750 | 0.927 |
| Ptpn5 | 0.04391 | 0.643 |
| Rps27 | 0.01114 | 0.608 |
| Psmb5 | 0.04123 | 0.500 |
| Fmnl1 | 0.02409 | 0.491 |
| Prmt1 | 0.03650 | 0.473 |
| Ndufaf4 | 0.00109 | 0.473 |
| Ddi2 | 0.03225 | 0.456 |
| Btf3 | 0.02712 | 0.434 |
| Spon1 | 0.02731 | 0.433 |
| Rab8a | 0.02410 | 0.397 |
| Eif1 | 0.01400 | 0.379 |
| G3bp2 | 0.02647 | 0.341 |
| Tpt1 | 0.00006 | 0.334 |
| Hsph1 | 0.00032 | 0.324 |
| Sh3kbp1 | 0.01395 | 0.291 |
| Rps16 | 0.00703 | 0.290 |
| Dnaja1 | 0.01025 | 0.280 |
| Grb2 | 0.01957 | 0.252 |
| Eef1a1 | 0.03566 | 0.235 |
| Eef2 | 0.00045 | 0.221 |
| Tra2a | 0.01602 | 0.215 |
| Rheb | 0.04696 | 0.214 |
| Kiaa0513 | 0.03774 | 0.206 |
| Rpl13 | 0.03803 | 0.205 |
| Hsp90ab1 | 0.02585 | 0.195 |
| Hpcal4 | 0.00062 | 0.191 |
| Hspa8 | 0.03850 | 0.131 |

Downregulated after KA

| Protein | Adj. p-value | Log2 FC (KA/Sal) |
|---|---|---|
| Bckdhb | 0.02584 | -0.810 |
| Tinagl1 | 0.04738 | -0.796 |
| Kcnj10 | 0.02556 | -0.758 |
| Naa15 | 0.00127 | -0.722 |
| Ank1 | 0.01913 | -0.649 |
| Abcd3 | 0.00731 | -0.624 |
| Snrpa1 | 0.01207 | -0.561 |
| Ggact | 0.02672 | -0.535 |
| Aldh4a1 | 0.02659 | -0.529 |
| Hectd4 | 0.00104 | -0.508 |
| Cygb | 0.02392 | -0.486 |
| Trappc8 | 0.02779 | -0.440 |
| Scg5 | 0.01695 | -0.430 |
| Ddhd1 | 0.01846 | -0.393 |
| Pnmal2 | 0.04917 | -0.356 |
| Akt1 | 0.00585 | -0.353 |
| C20orf27 | 0.02376 | -0.352 |
| Tiprl | 0.00950 | -0.337 |
| Erlin2 | 0.00276 | -0.300 |
| Hadha | 0.00548 | -0.300 |
| Sorbs1 | 0.04445 | -0.282 |
| Hadhb | 0.04115 | -0.277 |
| Alcam | 0.03304 | -0.265 |
| Scfd1 | 0.01639 | -0.265 |
| Pgls | 0.02616 | -0.262 |
| Hnrnpr | 0.02022 | -0.253 |
| Dgkg | 0.03988 | -0.247 |
| Agfg2 | 0.04260 | -0.244 |
| Ndufb5 | 0.03526 | -0.244 |
| Prxl2a | 0.01638 | -0.237 |
| Pitpnm1 | 0.04720 | -0.223 |
| Ahcy | 0.04338 | -0.184 |
| Atp5pd | 0.03795 | -0.174 |
| L1cam | 0.01534 | -0.167 |
| Actn1 | 0.01974 | -0.083 |

**B** **RGS14 KO**

Upregulated after KA

| Protein | Adj. p-value | Log2 FC (KA/Sal) |
|---|---|---|
| Spag7 | 0.00011 | 1.043 |
| Srrm2 | 0.01341 | 0.755 |
| Eif5b | 0.03604 | 0.736 |
| Cmip | 0.02918 | 0.733 |
| Mlst8 | 0.02592 | 0.705 |
| Dnajb5 | 0.04264 | 0.697 |
| Syt11 | 0.03861 | 0.626 |
| Sez6l | 0.00928 | 0.580 |
| Trappc8 | 0.03487 | 0.549 |
| Dlgap2 | 0.01553 | 0.495 |
| Cnn3 | 0.03324 | 0.477 |
| Scn1b | 0.00837 | 0.430 |
| Msi2 | 0.03794 | 0.407 |
| Kif3a | 0.02329 | 0.374 |
| Tbca | 0.03857 | 0.352 |
| Celf2 | 0.03192 | 0.326 |
| Sgta | 0.00067 | 0.292 |
| Ssr1 | 0.02485 | 0.289 |
| Asb8 | 0.04264 | 0.277 |
| Ggt7 | 0.03534 | 0.276 |
| Slc27a4 | 0.01662 | 0.276 |
| Farsb | 0.02048 | 0.257 |
| Tpt1 | 0.00416 | 0.237 |
| Rps9 | 0.04456 | 0.237 |
| Hsph1 | 0.01982 | 0.218 |
| P4hb | 0.02167 | 0.189 |
| Eef2 | 0.00978 | 0.171 |

Downregulated after KA

| Protein | Adj. p-value | Log2 FC (KA/Sal) |
|---|---|---|
| Ehbp1 | 0.02550 | -1.389 |
| Rnf141 | 0.04137 | -0.896 |
| Cacna1e | 0.00274 | -0.593 |
| Ptgr2 | 0.01712 | -0.461 |
| Srprb | 0.00560 | -0.457 |
| Mark3 | 0.01132 | -0.422 |
| Pmm1 | 0.00377 | -0.402 |
| Pip5k1a | 0.01768 | -0.397 |
| Rgs7bp | 0.00735 | -0.391 |
| Fth1 | 0.00248 | -0.371 |
| Mtmr1 | 0.01415 | -0.363 |
| Apoe | 0.01321 | -0.359 |
| Hagh | 0.03390 | -0.324 |
| Ncam1 | 0.03203 | -0.318 |
| Sorbs1 | 0.04719 | -0.312 |
| Mblac2 | 0.02923 | -0.294 |
| Mpp3 | 0.02482 | -0.288 |
| Abcb1a | 0.03589 | -0.285 |
| Sh3kbp1 | 0.04989 | -0.264 |
| Scfd1 | 0.04290 | -0.253 |
| Xpnpep1 | 0.00664 | -0.240 |
| Sh3bgrl | 0.03115 | -0.168 |
| Actr1b | 0.03090 | -0.148 |

**Supplemental Table 2. Comprehensive list of differentially expressed proteins (DEPs) one day after KA-SE in WT or RGS14 KO hippocampi.** (A) List of upregulated (left) or downregulated (right) DEPs one day following KA-SE in WT hippocampi, their adjusted p-values, and the fold change of abundance ($\log_2$ FC (KA/SAL)). (B) List of upregulated (left) or downregulated (right) DEPs one day following KA-SE in RGS14 KO hippocampi, their adjusted p-values, and the fold change of abundance ($\log_2$ FC (KA/SAL)). FC, fold change.
